# Decoding Articulatory and Phonetic Features: Somatotopic and Sensorimotor Representations in Native and Non-native Consonant Perception

**DOI:** 10.64898/2026.03.06.709780

**Authors:** Tzuyi Tseng, Simon Thibault, Jennifer Krzonowski, Mélanie Canault, Alice C. Roy, Claudio Brozzoli, Véronique Boulenger

**Affiliations:** CNRS, Université Lyon 2, Dynamique du Langage, Lyon, France; Aix-Marseille Université, Marseille, France; Centre de Recherche en Psychologie et Neurosciences, CNRS UMR 7077, Marseille, France; Institute for Language, Communication and the Brain, Aix-en-Provence, France; Université Lyon 1, Villeurbanne, France; Integrative Multisensory Perception Action & Cognition Team (ImpAct), Centre de Recherche en Neurosciences de Lyon, INSERM U1028, CNRS UMR5292, Université Claude Bernard Lyon 1, Lyon, France

**Keywords:** Speech Perception, Motor Cortex, Somatotopy, Sensorimotor Processing, Phonetic Features, Non-native Phonemes, fMRI, MVPA, RSA

## Abstract

Speech perception relies on the integration of auditory and articulatory information, yet the precise role of motor regions remains debated. We combined fMRI with multivariate pattern analyses (MVPA) and behavioral measures to investigate cortical representations of native French and non-native Mandarin consonant perception under clear and noisy conditions. Cross-modal classification analysis showed that articulator-specific features of perceived degraded native bilabial and dental consonants are mapped somatotopically in right lip and tongue motor areas, which are also engaged in consonant production. These representations may support consonant categorization by compensating for degraded auditory input. Representational similarity analysis further revealed that a network encompassing bilateral temporal auditory and frontal motor regions encodes phonetic features, including place and manner of articulation, for both native and non-native consonants. Our findings demonstrate that speech perception is inherently sensorimotor and that motor regions support both low-level embodied articulator-specific mapping and higher-level phonetic feature representations.

## INTRODUCTION

Scholars have long debated whether speech perception primarily involves auditory-acoustic or articulatory-acoustic mechanisms. According to the auditory account, speech sounds are decoded from spectrotemporal cues in the acoustic signal, and are mainly processed within the auditory temporal cortices (Diehl et al., 2004; Hickok, 2014). In this view, the neural substrates of speech perception are considered functionally independent from speech production circuits. In contrast, the articulatory view ascribes a central role of speech production processes in perception, suggesting that phonetic information in the acoustic signal is perceived as the articulatory gestures that generate those speech sounds (motor theory of speech perception, Liberman et al., 1967; Liberman & Mattingly, 1985, 1989; direct realist theory of speech perception, Fowler, 1986). Within this framework, speech perception is therefore thought to rely, at least in part, on the brain motor regions underlying speech production.

Consistent with these theoretical accounts, neuroimaging studies report the activity of both auditory and motor cortical areas during speech perception. Functional magnetic resonance imaging (fMRI) converges in showing that this process primarily engages the auditory cortex, particularly the bilateral superior temporal gyrus and sulcus (STG and STS) (Arsenault & Buchsbaum, 2015; Bonte et al., 2014; Evans & Davis, 2015; Fedorenko et al., 2024). In parallel, speech perception has been shown to activate the bilateral motor and premotor (PMC) cortices (Callan et al., 2010; Skipper et al., 2017; Tseng et al., 2026 for a review; Wilson et al., 2004). Notably, passive perception of bilabial consonants (e.g., /b/) activates the left dorsal precentral gyrus associated with lip movements, whereas dental consonant perception (e.g., /d/) recruits its ventral part, which is responsible for tongue movements (Pulvermüller et al., 2006; see corroborating findings using transcranial magnetic stimulation (TMS) in D’Ausilio et al., 2009). Such somatotopically organized motor activation suggests that speech perception selectively engages articulator-specific representations involved in producing the perceived consonants. TMS studies provided converging evidence by showing that motor regions play a functional role in speech sound processing, as stimulating or transiently inhibiting the motor cortex facilitates or impedes consonant perception, respectively (Möttönen & Watkins, 2009; Murakami et al., 2015; Sato et al., 2009; Skipper et al., 2017 for a review; Smalle et al., 2015).

Contemporary neurolinguistic models therefore emphasize the sensorimotor nature of speech perception, proposing that it emerges from tightly coupled action-perception interactions and complementary contributions of auditory and motor systems (Pulvermüller & Fadiga, 2010, 2016; Schomers & Pulvermüller, 2016; Laurent et al., 2017). Within this framework, the fundamental units of speech are viewed as perceptuo-motor rather than exclusively auditory or articulatory, providing a common representational basis that enables parity between speaker and listener (Schwartz et al., 2008, 2012). More broadly, this perspective aligns with embodied theories of cognition, which propose that language processing is grounded in sensory and motor experience (Barsalou, 2008; Fischer & Zwaan, 2008; Foglia & Wilson, 2013; Pulvermüller & Fadiga, 2010, 2016).

Beyond these representations specific to the movement and position of the articulators, sensorimotor regions may also encode higher-level representations related to phonetic features. Phonetic features, such as place and manner of articulation, categorize physical or acoustic properties of speech sounds, which describe characteristics of sounds independently of their linguistic function. FMRI studies using multivariate pattern analyses (MVPA) on consonant perception show that neural activity patterns in distributed networks encompassing not only the auditory temporal, but also the pre- and postcentral gyri, enable the decoding of place (e.g., labials vs. alveolars) and manner (e.g., plosives vs. fricatives) of articulation (Archila-Meléndez et al., 2018; Correia et al., 2015; Du et al., 2014; Evans & Davis, 2015). These findings therefore support that the processing of phonetic features relies on close interactions between auditory and motor systems. Chevillet and colleagues (2013) reached a similar conclusion using an fMRI rapid adaptation paradigm, in which participants listened to paired speech stimuli extracted from a place-of-articulation continuum (/da/-/ɡa/). They showed that the PMC selectively responds to perceived phoneme categories – showing reduced activation for stimuli within the same category but increased activity when pairs cross the category boundary - whereas auditory temporal regions track continuous acoustic variations (e.g., sound frequencies) irrespective of category (see also Dole et al., 2022 for congruent findings on a vowel continuum). Despite this accumulating evidence, several fMRI studies using both univariate and multivariate pattern analyses have however failed to find motor activity during speech perception (Szenkovits et al., 2012) or to replicate somatotopic activity in lip and tongue motor regions during the perception of labial and dental/alveolar consonants (Arsenault & Buchsbaum, 2016; see Schomers & Pulvermüller, 2016 for discussion on methodological issues). The precise functional role of motor cortex activity in speech perception therefore remains an open question, particularly whether it contributes to speech decoding through somatotopically organized representations of the articulatory movements underlying speech sound production.

One key argument in favor of the motor system’s functional role is its contribution to speech decoding under challenging listening conditions, such as in background noise (e.g., D’Ausilio et al., 2012; Du et al., 2014; Murakami et al., 2011; Osnes et al., 2011; but see Perron et al., 2024) or during foreign language perception (e.g., Callan et al., 2004, 2014; Schmitz et al., 2019; Wilson & Iacoboni, 2006). In a series of TMS studies, Nuttall et al. (2016, 2017) demonstrated that the perception of distorted syllables, whether due to articulatory impediments or masking noise, evoked larger motor potentials than clear syllables, a finding also observed in relation to participants’ hearing abilities. Furthermore, motor activity to degraded speech may even outweigh the auditory contribution (Brisson & Tremblay, 2021). Indeed, TMS-induced improvement in the identification of syllables embedded in multi-talker babble was larger after stimulation of the left PMC than of the left posterior STS. This is consistent with fMRI MVPA data showing that for noise-masked phonemes, only the left dorsal PMC and primary motor cortex (M1) encode phonemic information, showing similar neural patterns for within-category (e.g., two exemplars of /ba/) compared to between-category phonemes (e.g., /ba/ vs. /da/) (Du et al., 2014). Non-native speech similarly challenges listeners by introducing systematic deviations from native phonemes, even when overall acoustic clarity is preserved (Best, 1994). Using fMRI univariate approaches, several studies have reported stronger activations in the bilateral STG and STS and/or (pre)motor cortex for non-native than for native phoneme perception (Callan et al., 2014; Wilson & Iacoboni, 2006; see also Schmitz et al., 2019 for TMS evidence). For instance, when identifying the English /ɹ/-/l/ contrast, Japanese native speakers showed enhanced activity in a broad network, including the bilateral PMC and the inferior frontal gyrus (IFG, Broca’s area), whose activation level directly correlated with behavioral accuracy (Callan et al., 2004, 2014). This increased involvement suggests that the motor system may compensate for the lack of established articulator-specific representations for non-native speech sounds. More generally, these findings support the idea that motor regions may facilitate the mapping between auditory and articulatory representations by instantiating internal forward models (Callan et al., 2004; Iacoboni, 2008; Wilson & Iacoboni, 2006; see also Rauschecker & Scott, 2009), especially when speech perception becomes effortful. Yet, although both the auditory and motor cortices appear to be activated during the perception of degraded native and intact non-native phonemes, it remains unclear whether this sensorimotor involvement operates similarly across different languages and listening situations.

Overall, fMRI evidence for the role of the motor system during speech perception remains limited and inconsistent. In particular, whether motor activity is somatotopically (i.e. articulator-specifically) elicited during native speech perception is still debated and this question remains largely unexplored for non-native speech. Moreover, it is unclear whether and how sensorimotor regions encode distinct phonetic features, and whether these higher-level feature-based representations are shared both within and across native and non-native speech sounds. The current fMRI study directly addresses these gaps by systematically investigating neural activity patterns in motor and auditory temporal regions at both the articulator-specific and phonetic-feature representational levels during the perception of native and non-native consonants under optimal and degraded conditions. Going beyond prior work that primarily focused on intact native speech, we employed a dual fine-grained multivariate approach to identify shared and condition-specific neural representations of consonants across languages and perceptual conditions. First, we assessed whether consonant perception engages somatotopically organized motor representations. Using cross-modal decoding analysis, we tested whether neural patterns associated with lip and tongue articulatory movements could predict those elicited by the perception of bilabial and dental consonants, respectively, and whether this articulator-specific mapping varies across language and/or listening conditions. According to the motor account, motor representations should be engaged during speech perception regardless of whether the consonants are native or non-native. In contrast, from an embodied-cognition perspective, speech processing is shaped by sensorimotor experience. The somatotopic decoding of consonants should therefore be successful in the native language but be weaker or absent for non-native consonants for which listeners lack articulatory experience. Furthermore, in line with previous evidence, motor decoding should be particularly evident under challenging listening conditions, where consonants are masked by noise. As a second step, we examined whether speech sound representations are organized according to phonetic features and encoded within both motor and auditory regions. Using representational similarity analyses (RSA; Kriegeskorte et al., 2008), we tested whether the similarity structure of observed neural patterns aligns with theoretically predicted patterns based on the phonetic features of the perceived consonants. Representations of phonetic features constitute a higher-order level of representation than those of individual articulators and articulatory movements. Indeed, these features generalize across speech sounds and languages by grouping them into broader, discrete classes based on shared production properties (e.g., place and manner of articulation). If speech representations are sensorimotor in nature, both auditory and motor regions should encode phonetic features independently of language, provided these features are shared between native and non-native consonants. By combining cross-modal classification and RSA, we therefore aim to provide a comprehensive account of speech perception that maps the transition from low-level, articulator-specific motor representations to higher-order phonetic feature organization.

## RESULTS

### Behavioral categorization performance

Thirty-two healthy, right-handed native French adults participated in a combined behavioral and fMRI study, which featured native (/p/, /t/, /ʃ/ and /ʁ/) and non-native (Mandarin) consonants (/pʰ/, /tʰ/, /ʂ/ and /x/) embedded in consonant-vowel (CV) syllables. Participants were instructed to listen to triplets of identical syllables (e.g., /pʰa pʰa pʰa/), under intact or noisy perceptual conditions, and to categorize the heard consonant as native or non-native. These consonants capture key distinctions across the native and non-native consonant inventories by systematically varying articulator-specific configurations alongside phonetic features, namely place and manner of articulation, as well as laryngeal features (voicing and aspiration), across languages. Place (e.g., bilabial for /p/ and dental for /t/) and manner (e.g., plosives /p/, /t/, /pʰ/, /tʰ/ and fricatives /ʃ/, /ʁ/, /ʂ/, /x/) of articulation define where and how the consonants are produced, respectively. Aspiration corresponds to a burst of air at consonant release (e.g., aspirated /pʰ/ and /tʰ/ compared with unaspirated /p/ and /t/), whereas voicing refers to the vibration of the vocal folds (e.g., voiced /ʁ/ versus voiceless /x/).

The behavioral results revealed that participants categorized intact consonants significantly better than noisy ones, and native consonants better than non-native ones (Figure 1). This was confirmed by significant main effects of noise (*F*(1, 23) = 27.87, *p* < .001), language (*F*(1, 23) = 8.51, *p* = .008) and consonant (*F*(3, 69) = 73.62, *p* < .001) in a repeated measures ANOVA. The noise × consonant (*F*(3, 69) = 4.83, *p* = .004), noise × language (*F*(1, 23) = 5.43, *p* = .029), consonant x language (*F*(3, 69) = 37.32, *p* < .001) and consonant x language x noise (*F*(3, 69) = 10.62, *p* < .001) interactions were also significant. Tukey’s HSD post-hoc tests showed that noise significantly decreased categorization accuracy for the native bilabial /p/ (*p* < .001), both relative to the intact perceptual condition and to its non-native aspirated counterpart (/pʰ/; *p* < .001). For the fricatives, participants performed significantly worse for the non-native retroflex /ʂ/ than for the native post-alveolar /ʃ/, regardless of the perceptual condition. As a matter of fact, this non-native fricative, whether intact or noisy, elicited significantly lower performance than all other consonants (*p*_s_ < .001).

**Figure 1.**
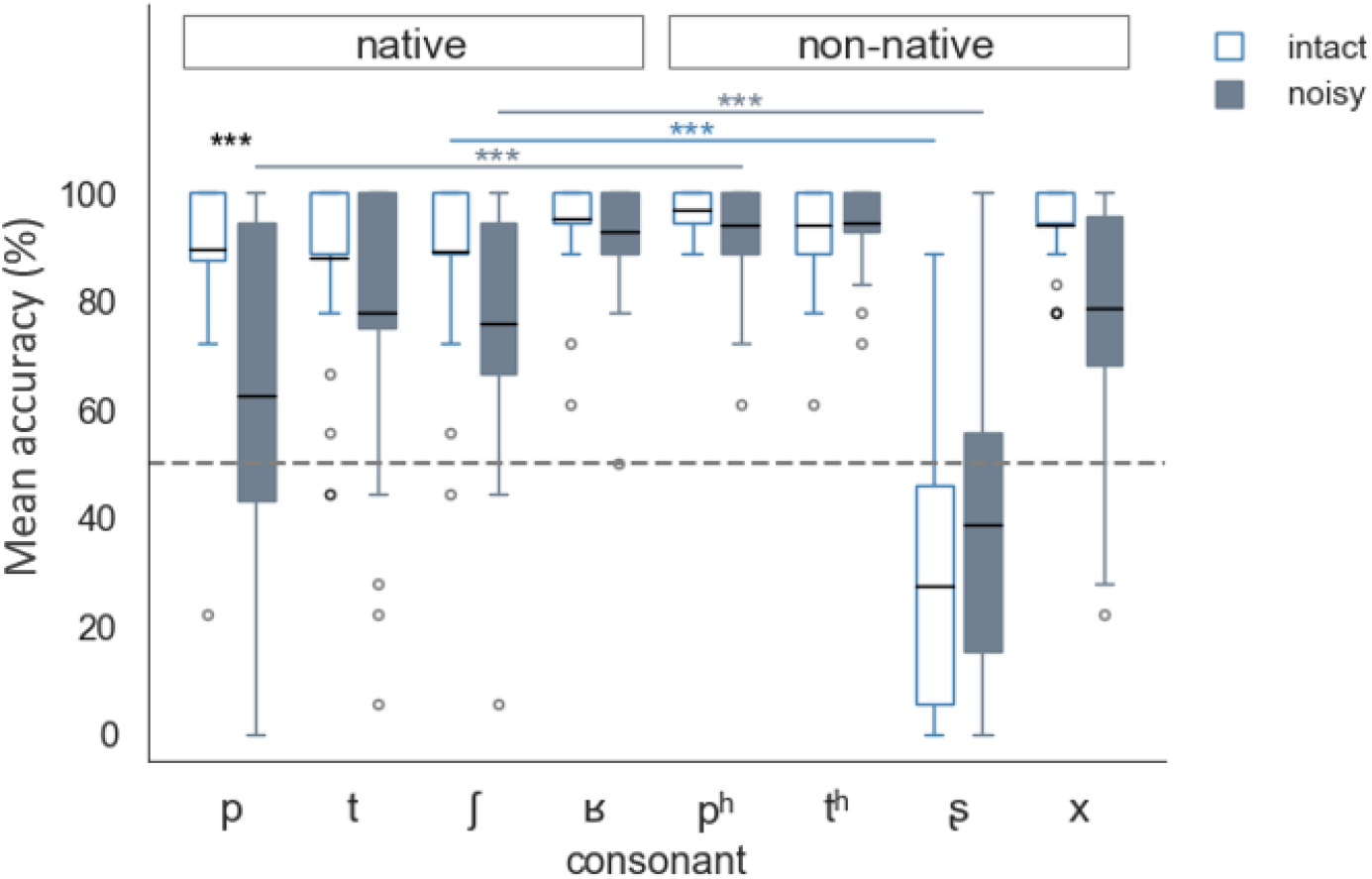
Behavioral categorization accuracy (mean % of correct responses) for native (/p/, /t/, /ʃ/, /ʁ/ on the left) and non-native (/pʰ/, /tʰ/, /ʂ/, /x/ on the right) consonants under intact (blue unfilled boxes) and noisy (grey filled boxes) conditions. Boxplots indicate the middle 50% (interquartile range) of the group accuracy scores in each condition, spanning from the first quartile (25th percentile) to the third quartile (75th percentile). The lines extending from each box capture data points outside this range, covering most of the remaining 25% on each side, with dots indicating outliers. The horizontal black line in each box represents the mean accuracy, and the dashed horizontal line shows chance level (50%). Asterisks illustrate significant differences between conditions (\*\*\**p* < .001): black asterisks indicate differences between intact and noisy conditions for each consonant, whereas blue and grey asterisks denote differences between native and non-native consonants (/p/-/pʰ/, /t/-/tʰ/, /ʃ/-/ʂ/ and /ʁ/-/x/) in the intact and noisy conditions, respectively.

### Articulator-specific patterns in the motor cortex during production predict patterns for consonant perception

Our univariate analyses showed robust activation of both auditory and (pre)motor cortices during consonant categorization across all language and perceptual conditions (see Supplemental information (SI), Figure S1 and Table S1). In particular, we found stronger right precentral activity for non-native compared to native consonants, irrespective of noise, whereas categorization of intact compared to noisy consonants elicited stronger activity in the STG, regardless of language (Figure S2). The corresponding reverse contrasts did not yield any significant clusters in either case. In addition, brain-behavior correlations showed that the better participants were at categorizing consonants in their native language, the stronger the right precentral activity (Figure S3A). Better non-native consonant categorization was instead associated with greater left inferior frontal activity (Figure S3B). Stronger activity in a different cluster of the left inferior frontal gyrus was also associated with better categorization performance under noisy conditions, regardless of language (Figure S3C).

We next assessed whether articulator-specific representations are somatotopically encoded in the motor cortex and recruited during consonant perception. Specifically, we tested whether neural activation patterns associated with articulatory movements predict those elicited during consonant perception. To this end, a classifier was trained to distinguish the neural patterns elicited by lip and tongue movements (i.e. within-modal classification) and then tested to classify those elicited by the perception of bilabial and dental consonants under intact and noisy conditions in the native and non-native languages (i.e. cross-modal classification). These classification analyses were performed within individual precentral regions-of-interest (ROIs) in both hemispheres defined from the articulatory localizer task (see *Methods*). We hypothesized that the classifier would successfully decode perception patterns for consonants produced with the corresponding articulators, especially under degraded listening conditions. In addition, while strict motor theories predict similar decoding performance across languages, the embodied-cognition account expects this cross-modal mapping to be weaker or absent for non-native consonants.

The within-modal classifier significantly discriminated the lip and tongue motor patterns within bilateral precentral ROIs (FDR-corrected *p* < .001, Cohen’s *d* = 3.61 in the left hemisphere and FDR-corrected *p* < .001, Cohen’s *d* = 3.52 in the right hemisphere), with high group mean accuracy (93%, SD = 12.04, 95% confidence interval (CI) [88.41, 97.83] in the left hemisphere and 95%, SD = 12.75, 95% CI [89.13, 99.28] in the right; Figure 2 left panel). Crucially, cross-modal classification revealed that neural patterns elicited by the perception of native bilabials and dentals (i.e. /p/ and /t/, respectively) in the noisy condition were successfully distinguished by the motor classifier within the right precentral ROI (mean accuracy 60%, SD = 13.7, 95% CI [54.18, 65.01], FDR-corrected *p* < .01, Cohen’s *d* = 0.69; Figure 2 right panel). Decoding performance did not significantly differ from chance in the other consonant conditions. To confirm the specificity of this effect, we conducted a control analysis using the same classifier to decode the neural patterns elicited by the perception of coronal (/ʃ/, /ʂ/) and dorsal fricatives (/ʁ/, /x/), which do not rely on corresponding lip and tongue articulatory features. As expected, classification performance did not significantly exceed chance for these fricatives (Figure S5), confirming that the observed cross-modal decoding reflects the activation of articulator-specific motor representations rather than general consonant discrimination.

**Figure 2.**
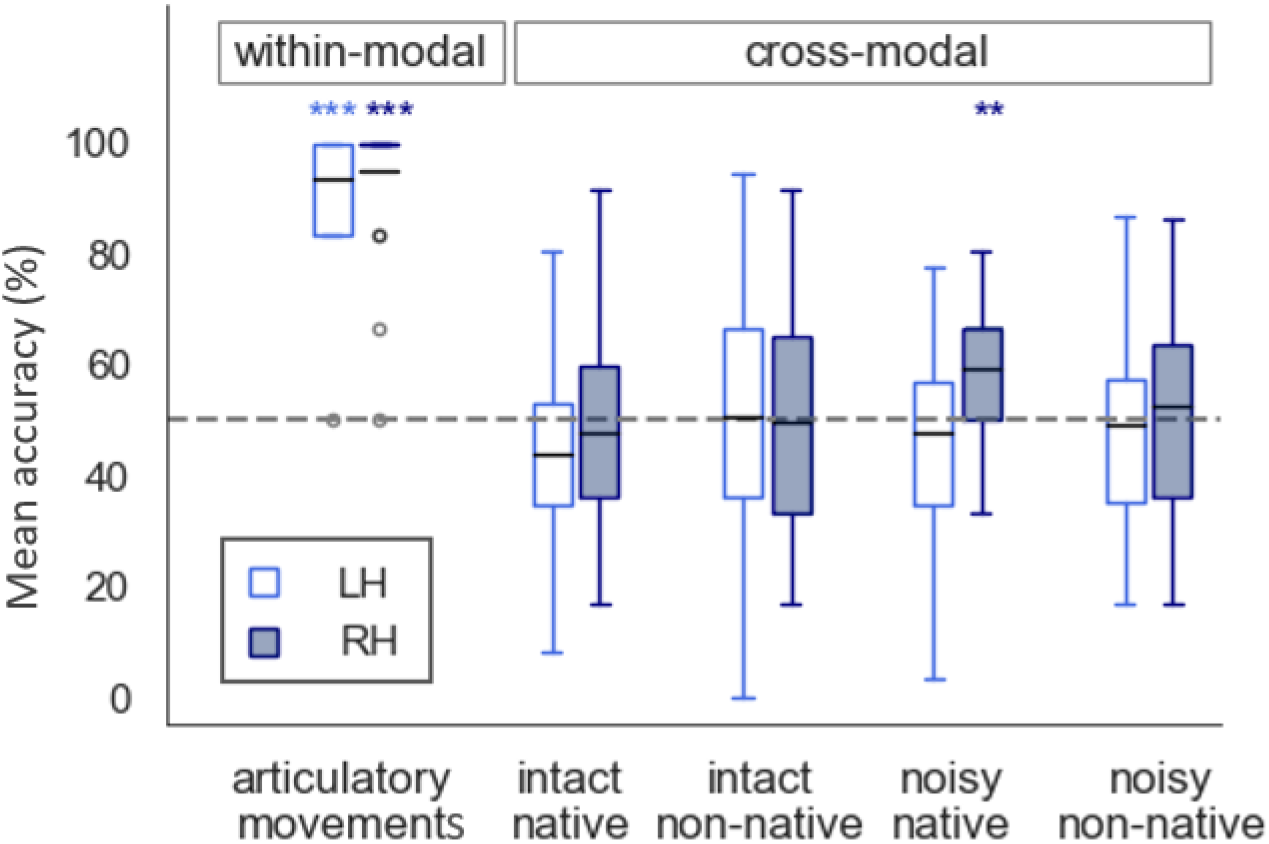
MVPA classification accuracy of motor activation patterns (within modality, first two blue boxplots on the left) for lip and tongue articulatory movements, and cross-modal pattern classification (the rest 8 boxplots on the right) for native and non-native bilabial and dental consonant perception in the intact and noisy conditions. For each condition, the first boxplot shows the mean classifier accuracy in the precentral ROI of the left hemisphere (LH, light blue unfilled boxes); the second one shows the accuracy in its counterpart in the right hemisphere (RH, dark blue filled boxes). The black horizontal line in each box indicates the group mean accuracy for the corresponding condition. Asterisks (\*\**p* < .01; \*\*\**p* < .001) indicate statistical significance in the one-sample permutation test against chance level (50%, dashed line) with FDR correction for multiple comparisons across the four conditions in the two hemispheres.

### Auditory and motor regions encode phonetic-feature representations

To further study the neural encoding of speech sounds at a higher representational level, namely whether and how perceived phonetic features are represented in sensorimotor cortices, we conducted representational similarity analyses (RSA) using a searchlight approach constrained to a set of seven predefined ROIs (see *Methods*; see Correia et al., 2015 for a similar approach). These ROIs (see shaded grey areas in Figure S6) are known to be involved in speech processing (Archila-Meléndez et al., 2018; Arsenault & Buchsbaum, 2015; Correia et al., 2015; Du et al., 2014; Evans & Davis, 2015), and were also activated by consonant perception in the present study, as shown by our univariate analyses (Figures S1, S2, S3). The aim of these RSA was to identify cortical regions where neural patterns discriminate the phonetic features of the perceived consonants, irrespective of language (native or non-native) and perceptual conditions (intact or noisy). In other words, we sought to detect regions where consonants sharing the same phonetic features elicit similar neural patterns. Neural representational dissimilarity matrices (RDMs) were computed to capture the pairwise correlations between the observed activity patterns associated with each perceived consonant. These empirical RDMs were then correlated with theoretical RDMs modelling the predicted within-feature similarity and between-feature dissimilarity. The predicted similarity structures were modelled based on the four following phonetic features: place of articulation (Figure 3A), manner of articulation (Figure 3B), aspiration (Figure 3C) and voicing (Figure 3D; see *Methods* and also Figure S7A for an additional theoretical model of language identity). A high correlation between observed neural and predicted RDMs indicates that neural activity patterns within a given cortical region encode the phonetic feature of interest.

**Figure 3.**
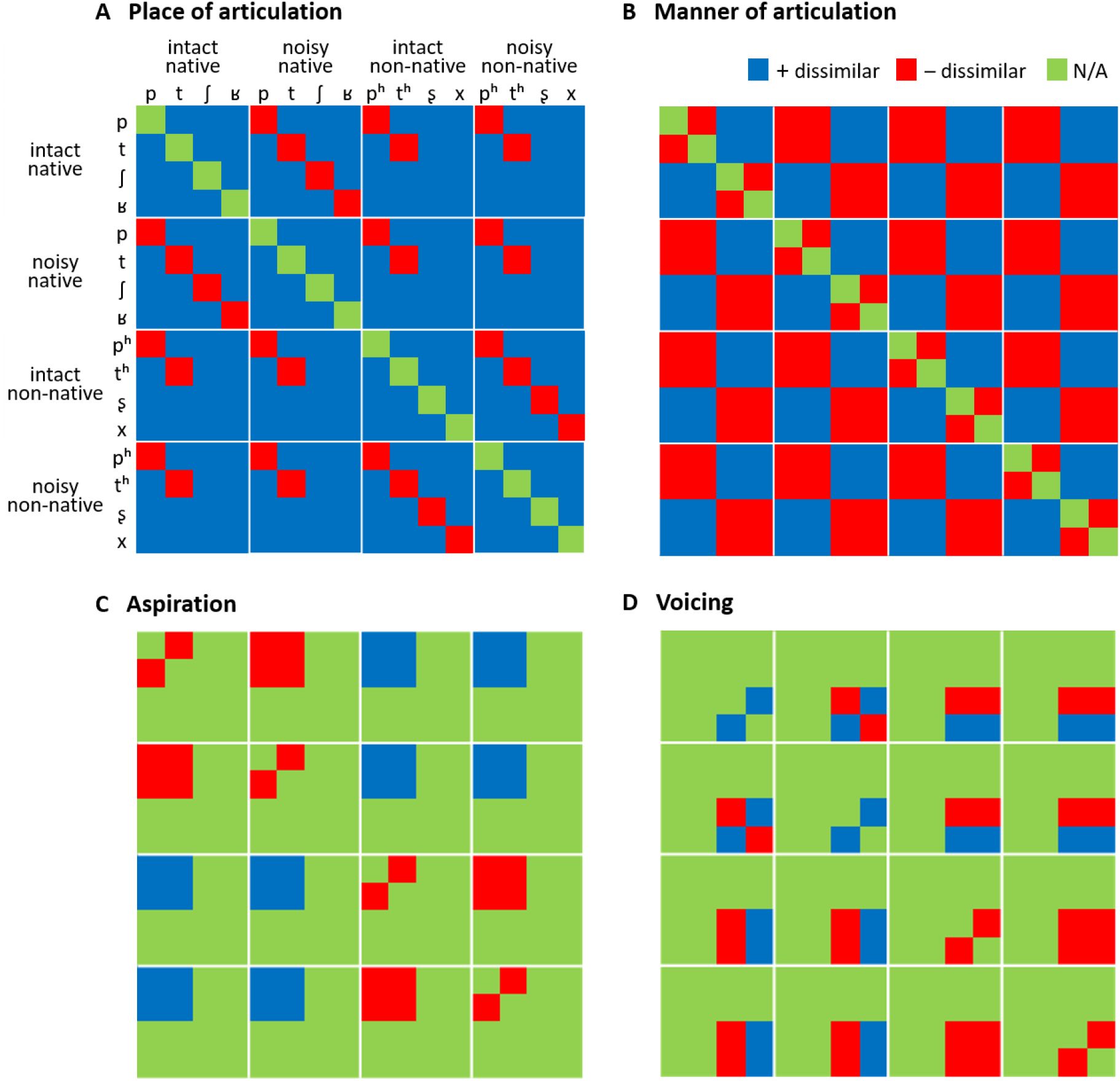
Theoretical feature-based RDMs predicting the similarity structure of the neural patterns for (A) place of articulation, (B) manner of articulation, (C) aspiration and (D) voicing. Each tile represents the hypothesized similarity/dissimilarity between the neural activity patterns for two consonants, in the intact native, noisy native, intact non-native and noisy non-native conditions. Red tiles express increased similarity (with a value of -1, i.e. decreased dissimilarity); blue tiles express decreased similarity (with a value of 1, i.e. increased dissimilarity); green tiles express non-relevant associations (with N/A values, see Evans & Davis, 2015).

As shown in Figure 4, consonants sharing the same place of articulation (yellow clusters) elicited similar neural patterns within a distributed bilateral network encompassing the pre- and postcentral gyri, insula, STG and MTG as well as the right IFG. Encoding of these features was more focal in the left hemisphere, particularly within dorsal motor and superior temporal areas, while it was more widely distributed across the right sensorimotor regions (see also Figure S6A and Table S2). For manner of articulation (green clusters in Figure 4; Figure S6B and Table S2), clusters were distributed bilaterally in a network largely overlapping with that identified for place of articulation and including the dorsal precentral gyri, IFG, insula, STG and MTG. Compared with place of articulation, manner of articulation was associated with stronger and more focal involvement of the bilateral insula and the right anterior superior temporal cortex, alongside a notable extension to the left anterior and posterior temporal cortex compared to place of articulation. Encoding of aspiration was more spatially restricted, with a single cluster in the right dorsal precentral gyrus (red cluster in Figure 4; Figure S6C and Table S2). Finally, clusters encoding voicing were primarily located in the left hemisphere, covering the ventral precentral gyrus and IFG, as well as the bilateral STG and the right posterior MTG (purple clusters in Figure 4; Figure S6D and Table S2). Overall, these findings show that during consonant perception, the similarity structure of phonetic features is encoded within a broad network spanning both motor and auditory regions in the left and/or right hemispheres. Crucially, the precentral gyrus and the superior and middle temporal gyri are consistently involved across languages and perceptual conditions. A supplementary analysis showed that consonants belonging to the same language (native or non-native) elicited activity in a network distributed over bilateral pre- and post-central gyri, as well as the right supramarginal, superior and middle temporal gyri (Figure S7B).

**Figure 4.**
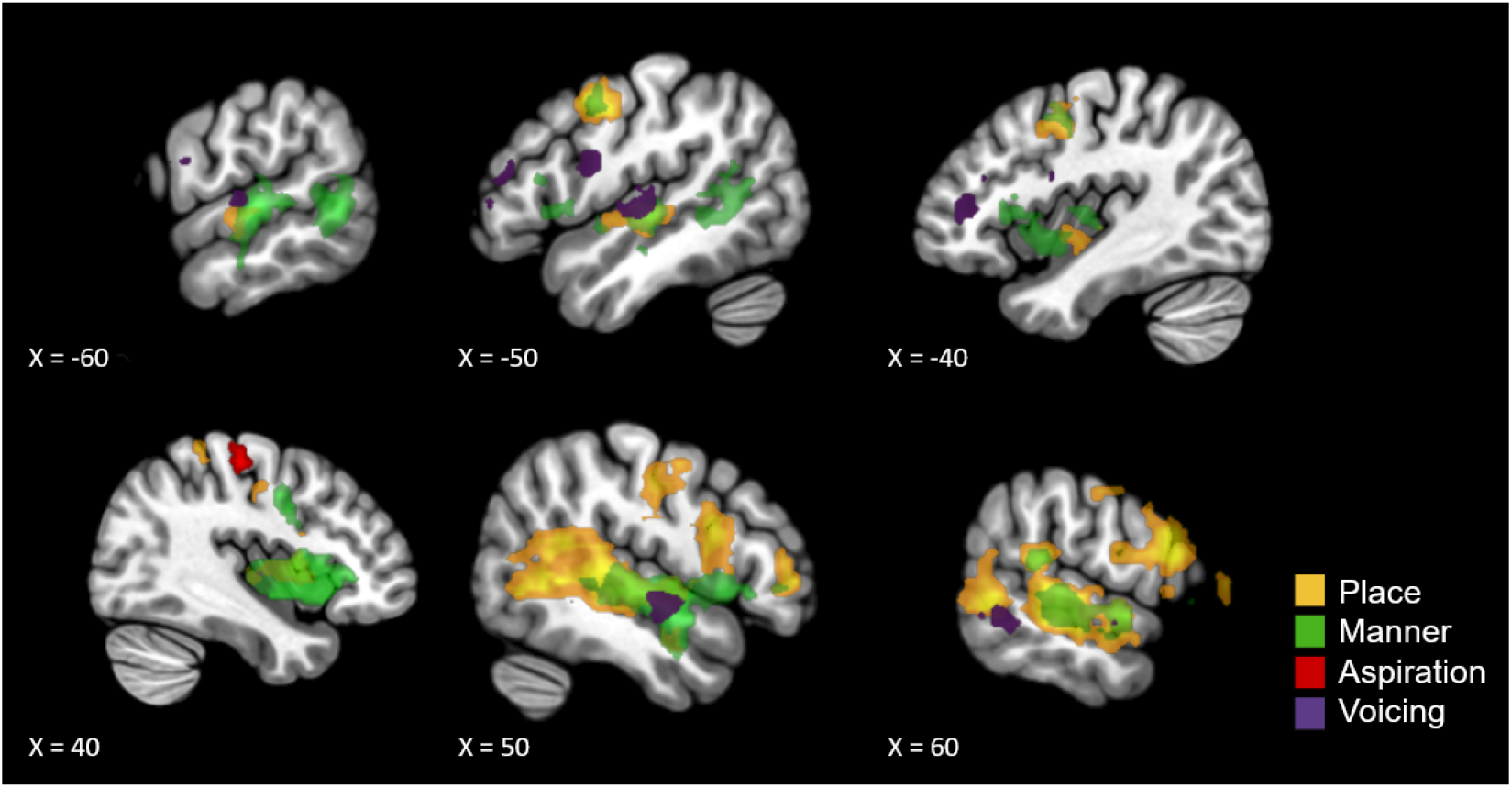
Searchlight RSA brain maps displaying the overlap of neural activity patterns associated with the four phonetic features of interest during consonant perception. Brain maps show the significant clusters after permutation testing thresholded at *p* < .05 (FDR-corrected) with a minimum cluster size of 780 mm^3^. Colors indicate the examined phonetic features: yellow for place of articulation, green for manner of articulation, red for aspiration and purple for voicing.

To better characterize the neural representations of these phonetic features, we then tested whether they were more strongly encoded in sensory, motor, or sensorimotor regions, and whether this encoding showed hemispheric dominance. Specifically, to assess the relative contributions of bilateral auditory and motor regions to phonetic feature encoding, we conducted a repeated-measures ANOVA on Spearman’s rho values extracted from the voxels in the left and right STG and precentral gyri that showed significantly similar neural patterns for consonants sharing the same phonetic features. Because significant RSA patterns for aspiration (red cluster in Figure 4) and voicing (purple clusters in Figure 4) were restricted to a single hemisphere or ROI (e.g., aspiration was encoded only in the right precentral gyrus), these features were excluded from the ANOVA. This analysis thus included rho values for place and manner of articulation only, which showed significant RSA patterns across all ROIs in both hemispheres. Three factors were included: Phonetic Feature (place vs. manner), ROI (STG vs. precentral gyri), and Hemisphere (left vs. right), allowing us to test whether representational similarity differed across cortical regions and hemispheres.

The repeated-measures ANOVA revealed significant main effects of Hemisphere (*F*(1,23) = 9.94, *p* = .004) and ROI (*F*(1,23) = 12.06, *p* = .002), as well as a significant Hemisphere x ROI interaction (*F*(1,23) = 10.48, *p* = .004). Post-hoc paired t-tests comparing the correlation values in the two hemispheres within each ROI, showed significantly higher values in the right than in the left STG (*t*(23) = -3.57, *p* = .002). This indicates that right temporal regions were more engaged in processing place and manner of articulation than their left counterparts. No such inter-hemispheric difference was observed within the precentral gyri. We then compared correlation values between the two ROIs within each hemisphere. Only the right hemisphere showed significantly greater values in the STG than in the precentral gyrus (*t*(23) = -3.15, *p* = .005). This suggests that the right superior temporal cortex encoded the two phonetic features more robustly than the right motor cortex. In contrast, no significant difference was observed between STG and precentral gyrus in the left hemisphere.

In summary, the RSA results reveal that the representations of perceived phonetic features that distinguish consonants, both within and across languages, are primarily encoded in a bilateral network of sensory and motor regions. Notably, features such as place and manner of articulation were found to be represented to a similar extent in auditory and motor cortices in the left hemisphere, whereas in the right hemisphere, encoding showed an auditory advantage.

## DISCUSSION

In this study, we combined behavioral and fMRI approaches to investigate the neural coding of perceived consonants in native and non-native languages, under both clear and degraded listening conditions. A key finding is that the right motor cortex maps articulator-specific representations of degraded native speech sounds in a somatotopic manner. Motor activation patterns associated with lip and tongue articulatory movements mirror those elicited by the perception of noise-masked native bilabial and dental consonants, respectively. Moreover, using RSA, we show that the representations of phonetic features distinguishing perceived consonants - both within and across native and non-native languages - are encoded in a distributed network encompassing bilateral motor and auditory cortices, supporting the view that both systems jointly contribute to speech perception.

Our fMRI univariate analyses first confirmed that both auditory and motor regions are engaged during speech perception. Importantly, motor cortical activity varied with behavioral performance: better categorization accuracy for native consonants was associated with stronger precentral activation. These findings support the idea that accurate phoneme perception benefits from the recruitment of articulatory representations in the motor cortex, echoing previous evidence for the functional role of motor regions in successful phoneme identification (e.g., Callan et al., 2010; Alho et al., 2014). However, enhanced motor activity did not systematically translate into better behavioral performance. Motor activity was indeed stronger for non-native than for native consonants, despite overall lower categorization accuracy for the former. This pattern of performance was especially observed for the retroflex fricative /ʂ/, which is not part of the native (French) phonological inventory. According to the Perceptual Assimilation Model (PAM, Best, 1994; Best et al., 2001), such difficulties arise when non-native contrasts rely on articulatory features that are not represented in the listener’s native repertoire, as is the case here for tongue retroflection. Interestingly, participants were able to accurately categorize non-native aspirated consonants (i.e. /pʰ/ and /tʰ/), even though aspiration does not constitute a phonemic contrast with their unaspirated counterparts in French (i.e. /p/ and /t/). Under degraded conditions, aspirated (non-native) plosives were even categorized more accurately than unaspirated (native) consonants (see Liang et al., 2023 for converging findings). This may be explained by the greater perceptual salience of voice-onset-time (VOT) cues for aspirated compared to unaspirated consonants (Table S3), which likely facilitates discrimination between the two, especially when consonants share the same place of articulation (i.e. bilabial for /p/ and /pʰ/, and dental for /t/ and /tʰ/ here). Taken together, these findings suggest a dissociation between native and non-native speech processing in the motor cortex (see Figure S7 for corroborating findings using an additional RSA on language identity).

While native consonants show a close correspondence between accuracy and motor involvement, stronger motor activity for non-native consonants is not associated with better behavioral performance. Such enhanced motor activity may reflect the engagement of internal forward models attempting, unsuccessfully, to simulate articulatory gestures that are absent from the listener’s native motor repertoire (Callan et al., 2004; Rauschecker & Scott, 2009; Wilson & Iacoboni, 2006). The successful categorization of non-native consonants may instead rely on the left IFG, as indicated by the significant correlation between activity in this region and behavioral accuracy (Figure S3B, MNI peak [-48, 10, 30]). This interpretation aligns with a recent coordinate-based meta-analysis which demonstrated a functional segregation within the speech production network during speech perception under challenging listening conditions (Perron et al., 2024; see also Alain et al., 2018). Specifically, Perron and colleagues showed that while the left IFG is engaged during speech-in-noise processing (MNI peak [-48, 8, 26]), this region exhibits an overlap with speech production only for linguistically complex speech (MNI peak [-54, 8, 8]). This dissociation led the authors to propose that the left IFG operates in a context-dependent manner, being more sensitive to linguistic variations than to mere signal quality. Although their meta-analysis mostly focused on native speech, our results extend this framework to non-native consonant categorization, by showing that the left IFG is specifically recruited when processing challenging linguistic input.

To summarize, our univariate results revealed a robust main effect of language, characterized by stronger motor activity for non-native compared to native consonants. Crucially, when considering categorization performance, we observed distinct regional profiles: greater precentral activity was associated with better native consonant categorization, whereas greater inferior frontal activity was associated with better performance under adverse conditions. Building on these univariate and behavioral findings, we next discuss how our MVPA findings allow to untangle the articulator-specific and phonetic-feature representation levels of speech sounds across language and listening conditions. These results further inform current theories of speech perception and provide insights into the embodied and cross-linguistic processing of perceived speech sounds.

### Somatotopic organization of articulator-specific motor representations during speech perception

Whether motor regions are somatotopically activated during consonant perception as a function of their articulator-specific representations remains unclear, with mixed results in the literature (Arsenault & Buchsbaum, 2016; Pulvermüller et al., 2006; Schomers & Pulvermüller, 2016; Szenkovits et al., 2012). Here, we provide new evidence that articulator-specific features of perceived consonants are mapped onto their corresponding representations within the motor cortex. Using a multivariate approach, a cross-modal classifier that was trained on motor activation patterns elicited by articulatory lip and tongue movements successfully decoded neural activity in the right precentral gyrus evoked by the perception of degraded native lip-(labial) and tongue-articulated (dental) plosive consonants, respectively. This finding demonstrates somatotopic mapping of articulator-specific features across speech production and perception, in line with Pulvermüller et al.’s findings (2006). Their univariate fMRI results indeed revealed a double dissociation within motor articulatory regions during consonant perception: passive listening to dental consonants preferentially activated the tongue motor area over the lip area, while the reverse was observed for labial consonants (see also Dmitrieva et al., 2025 for converging findings during intact word comprehension). Our multivariate results offer further support for this somatotopic organization by demonstrating that right motor regions underlying the articulatory movements involved in consonant production can decode the neural patterns elicited by consonant perception in noise. Crucially, this effect was selective, depending on both the articulator-specific features of the perceived consonants and the listeners’ language experience. First, decoding succeeded for bilabial and dental plosives (/p/ and /t/), which involve a complete closure of the vocal tract and clearly distinct articulators (lips vs. tongue, respectively), but failed for coronal and dorsal fricatives (/ʃ/ and /ʁ/), which rely on lingual constriction without full closure. This failure likely stems from the fact that fricatives involve articulatory configurations that do not align well with those used to train the classifier (i.e. bilabial /m/ and dental /l/). Indeed, while dental plosives (/t/) share a close articulatory configuration with the produced approximant /l/, fricatives require a more posterior and constricted tongue shape, thereby reducing the motor overlap with this consonant. This structural proximity may explain why successful cross-modal decoding was restricted to plosives. Our findings therefore indicate that articulator-specific representations are recruited during speech perception in an effector-specific manner, and that this production-perception overlap is constrained by precise articulatory correspondence. Second, cross-modal decoding succeeded only for noisy native consonants. This finding highlights two key points: motor cortical engagement during speech perception is specifically enhanced under challenging perceptual conditions and it is shaped by language-related sensorimotor experience. Successful classification of native consonants in the presence of noise but not in clear conditions aligns with previous evidence that motor activity likely supplements impoverished auditory input by retrieving articulatory representations from the motor repertoire (Callan et al., 2004; D’Ausilio et al., 2009, 2012; Du et al., 2014; Evans & Davis, 2015; Rauschecker & Scott, 2009), thereby supporting speech decoding through shared neural circuits for perception and production (Dmitrieva et al., 2025; Pulvermüller & Fadiga, 2010, 2016; Pulvermüller et al., 2006). Conversely, unsuccessful decoding for intact native consonants may reflect weaker and less consistent motor engagement when speech identification is relatively easy (see D’Ausilio et al., 2012 for congruent findings). This interpretation is consistent with our univariate analyses, which revealed greater activity in bilateral superior temporal areas when consonant categorization occurred under easier perceptual conditions (Intact vs. Noisy; Figure S2B). Taken together, these findings reinforce the idea that temporal auditory processing may predominate when the speech signal is clear, whereas motor contributions become increasingly important for speech decoding under challenging listening conditions (see also Laurent et al., 2017; Moulin-Frier et al., 2012). The successful classification for native consonants but not for non-native ones furthermore points to the critical role of embodied representations in speech processing (Pulvermüller & Fadiga, 2010, 2016; Pulvermüller et al., 2006). During native language acquisition, repetitive coupling between perception and production of speech sounds shapes fine-grained sensorimotor mappings that are grounded in the neural circuitry for language. In contrast, listeners lack motor expertise with non-native speech sounds, meaning these established mappings are absent. Accordingly, in the present study, the perception of non-native consonants failed to evoke precise, overlearned articulator-specific representations, preventing successful cross-modal decoding of the associated neural patterns.

Altogether, our cross-modal decoding findings support a functional role of the motor system in speech perception, in line with embodied cognition frameworks (Pulvermüller & Fadiga, 2010, 2016) and models based on forward internal predictions (Callan et al., 2004; Rauschecker & Scott, 2009). Specifically, we show that not only are motor regions activated during mere consonant perception but that this motor involvement is fundamentally shaped by the listener’s sensorimotor experience. We propose that these embodied articulator-specific representations may contribute to resolving degraded or ambiguous speech signals.

### Encoding of phonetic-feature representations of perceived consonants in auditory and motor regions

We further examined a higher representational level of speech sounds by investigating how phonetic features, namely place and manner of articulation, aspiration, and voicing, are represented in speech-related brain regions. Using an RSA approach, we captured the neural patterns reflecting the modelled similarity structures of these phonetic features during the perception of native and non-native consonants. These phonetic feature-based representations were found to be mapped onto a cortical network encompassing bilateral temporal and frontal motor regions, for both languages and under both clear and degraded listening conditions. This widespread distribution strongly supports a sensorimotor account of speech processing, demonstrating that when categorizing heard consonants, listeners access phonetic representations that capture both sensory and motor properties of the speech sounds. Crucially, this distributed encoding operates irrespective of language identity, provided that the underlying phonetic features are shared between the native and non-native inventories.

Neural patterns in bilateral precentral and superior temporal regions encoded the modelled place-of-articulation feature: these regions showed similar neural activity for native and non-native consonants sharing the same place of articulation (e.g., bilabial for intact /p/ and /pʰ/, uvular for intact and noisy /ʁ/). These patterns were more focal in the left hemisphere and more distributed in the right. Similarly, the perception of consonants sharing the same manner of articulation (e.g., plosive for intact /t/ and noisy /pʰ/, fricative for intact /ʂ/ and intact /ʁ/) induced focal bilateral precentral activity alongside more distributed bilateral superior temporal activation. This extensive bilateral sensorimotor engagement likely reflects the flexible perceptual mechanisms required to decode these inherently sensory and motoric features (Correia et al., 2015; Zheng et al., 2025). Specifically, the perceptual decoding of place and manner of articulation may rely on phonetic-feature representations that categorize shared properties of dynamically coordinated articulatory gestures involving multiple articulators, such as lips, tongue, jaw, and associated articulatory structures during production (Perrier, 2006). Notably, the broader superior temporal clusters observed for manner of articulation may reflect the fact that this feature requires the extraction of additional temporal cues, such as the degree and duration of articulatory closure, which contribute to manner distinctions (Fuchs & Birkholz, 2019). Consistent with this interpretation, direct cortical recordings in the STG showed that auditory responses are more strongly organized by manner than by place of articulation, reflecting high sensitivity to the degree rather than the location of articulatory constriction in the vocal tract (Mesgarani et al., 2014).

In light of our findings of bilateral feature-based representations, and given previous reports of hemispheric and regional differences in phonetic encoding, we further examined whether STG and precentral gyri encode place and manner of articulation to the same extent across hemispheres. In the left hemisphere, both regions contributed equally. In the right hemisphere, however, STG exhibited significantly stronger encoding than both the ipsilateral precentral gyrus and the left STG (consistent with Arsenault & Buchsbaum, 2015, but see Archila-Meléndez et al., 2018). These findings not only corroborate the intrinsic sensorimotor nature of speech sounds (Laurent et al., 2017; Schomers & Pulvermüller, 2016; Schwartz et al., 2008, 2012), but also highlight the functional role of both bilateral motor and auditory systems in representing phonetic features, with a potential right-hemispheric advantage. Such a contribution of the right hemisphere may support enhanced speech decoding across languages and perceptual conditions, broadening the scope of previous studies that restricted their investigations of these phonetic features to intact native speech (Archila-Meléndez et al., 2018; Arsenault & Buchsbaum, 2015; Correia et al., 2015).

In contrast, perceiving consonants sharing aspiration or voicing features did not engage bilateral auditory and motor regions as extensively as place and manner of articulation. These phonetic features primarily categorize the properties of speech production that depend on laryngeal opening and its temporal coordination with oral gestures (Fuchs & Birkholz, 2019; Jackson, 2000; Löfqvist, 1995). A cluster in the right dorsal precentral gyrus encoded the modelled aspiration feature, showing similar neural activity across perceived aspirated plosives (e.g., aspiration in intact or noisy /pʰ/ and /tʰ/). Conversely, voicing in fricatives (e.g., intact and noisy voiced /ʁ/, or intact and noisy unvoiced /x/) was encoded focally in left motor regions and bilateral superior temporal regions. Unlike the analyses for place and manner that included all consonants, RSA for aspiration were restricted to plosives and RSA for voicing to fricatives, which may account for the more localized activation and reduced statistical power observed. Beyond these methodological considerations, the right dorsal precentral cluster associated with the encoding of aspiration (MNI peak [38, -19, 56]) aligns with prior evidence implicating this region in fine laryngeal motor control (MNI peak [43, -18, 38], Eichert et al., 2020; [40, -5, 50], Liang et al., 2023). Such laryngeal control is crucial for regulating voice-onset time (VOT), which constitutes a distinctive feature between aspirated and unaspirated consonants, with typically longer values for the former. Our findings are consistent with fMRI studies showing that this dorsal precentral region contributes to detecting subtle VOT differences in plosives (Liang et al., 2023). Voicing, on the other hand, primarily reflects the presence or absence of vocal fold vibration. Specifically, voiced fricatives require precise laryngeal-oral coordination, with a partially closed glottis sustaining phonation while allowing airflow for oral turbulence (Fuchs & Birkholz, 2019). Perception of this feature likely involves the activation of neural representations for both motor commands and ongoing auditory monitoring of acoustic outcomes. Accordingly, in our study, voicing was encoded in a separate, more inferior left precentral cluster (MNI peak [-51, 2, 24]), lying relatively close to the ventral part of the larynx motor area (MNI peak [-58, -2, 18], Eichert et al., 2020; [-58, 3, 25], Tamura et al., 2022). This region is involved in the production of vocalizations and voiced consonants (Bouchard et al., 2013; Grabski et al., 2012; Dichter et al., 2018; Eichert et al., 2020; Toyoda et al., 2014) as well as in the perception of voicing (Tamura et al., 2022). Notably, we also found encoding of voicing in bilateral superior temporal regions, in line with previous MVPA studies that reported a predominant involvement of auditory regions for this feature (Correia et al., 2015).

### The inherent sensorimotor nature of speech sounds

Our RSA findings therefore point to a contribution of sensorimotor brain regions to the representation of phonetic features during speech perception. However, the question remains whether speech sounds are primarily stored as sensory, motor or integrated sensorimotor representations in the brain. First, purely auditory accounts suggest that speech perception relies on acoustic feature analysis within the temporal auditory cortex, viewing motor cortical activation as epiphenomenal (Diehl et al., 2004; Hickok, 2014). For instance, intracranial recordings in the motor cortex revealed that neural responses during speech perception reflected the acoustic properties of speech sounds, closely resembling responses observed in the auditory cortex (Cheung et al., 2016). This finding has been taken to suggest that the motor cortex contributes to speech perception by processing auditory rather than articulatory information, thereby indicating predominantly sensory-based representations. Our data clearly challenge this view, as both the successful cross-modal decoding in the motor cortex, reflecting the activation of articulator-specific representations, and the distributed encoding of phonetic features in frontotemporal networks indicate that motor regions are actively and specifically engaged in consonant perception, particularly under adverse listening conditions. Second, motor frameworks, such as the motor theory of speech perception (Liberman et al., 1967; Liberman & Mattingly, 1985, 1989) and the direct realistic theory of perception (Fowler, 1986), propose that the fundamental units of speech are inherently articulatory gestures. The frontal motor cortex is therefore suggested to play a dominant role in speech perception. Supporting this, recent fMRI MVPA has shown that neural patterns in motor regions generalized across perceived consonants sharing the same place of articulation, consistent with the retrieval of motor-based, articulatory representations (Archila-Meléndez et al., 2018). By demonstrating univariate motor activity to speech sounds, somatotopic mapping between perception and production, and encoding of phonetic features such as place of articulation in precentral areas, our results support these motor views, however only partially. Indeed, the failure to decode fricatives alongside the asymmetries observed between native and non-native languages in our classification analyses demonstrate that this motor mapping is not invariant. Instead, it is highly constrained by the listener’s own articulatory experience with the perceived speech sounds, depending on the precise articulatory alignment between the heard consonants and the listener’s motor repertoire.

Altogether, our findings are more consistent with contemporary sensorimotor accounts (Pulvermüller & Fadiga, 2010, 2016; Schomers & Pulvermüller, 2016; Laurent et al., 2017). Previous studies supporting this view have shown that the representations of place of articulation were encoded in the right precentral and left superior temporal gyri, whereas those of manner of articulation, thought to be more representative of high-order acoustic features, were encoded in left IFG and STG regions (Zheng et al., 2025; see also Cheung et al., 2016; Mesgarani et al., 2014). By demonstrating that phonetic features are encoded in both auditory and motor cortices, our RSA results support a dual auditory-motor integration framework (see also Correia et al., 2015). Rather than relying on a single motor module or on purely auditory analysis, speech perception involves tightly coupled sensorimotor circuits where articulator-specific motor representations interact with higher-level phonetic features to optimize speech processing across varying perceptual demands.

### The role of the right motor cortex in decoding perceived speech sounds

Despite the frequently reported left-lateralized involvement of motor regions in speech sound processing (e.g., Callan et al., 2010; D’Ausilio et al., 2009, 2012; Evans & Davis, 2015; Möttönen & Watkins, 2009), our findings point to a substantial contribution of right-hemisphere motor areas. This aligns with previous neuroimaging work showing either bilateral or right-dominant motor activity during speech perception and production (e.g., Behroozmand et al., 2015; Cogan et al., 2014; Dole et al., 2022; Preisig et al., 2022; Sheng et al., 2019; see Skipper et al., 2017 and Tseng et al., 2026 for reviews). Specifically, our cross-modal MVPA revealed somatotopic mapping of perceived degraded native consonants onto articulator-specific patterns restricted to the right motor cortex. In addition, when comparing motor engagement across hemispheres, we found no left hemispheric advantage in the encoding of phonetic features, place and manner of articulation in particular. Recent fMRI studies employing MVPA across both hemispheres similarly lack consistent evidence for left-hemisphere dominance. Instead, sensorimotor representations of phonetic features during intact native speech perception have more frequently been reported in the right precentral gyrus (Archila-Meléndez et al., 2018; Correia et al., 2015; Zheng et al., 2025) rather than exclusively in the left (Du et al., 2014).

Studies systematically examining motor involvement across both hemispheres remain comparatively rare. Among fMRI investigations using univariate analyses, precentral activation has been reported in the two hemispheres during both degraded native speech (e.g., Callan et al., 2010) and intact non-native speech perception (e.g., Callan et al., 2014). Notably, a right motor contribution has been reported under increased perceptual demands, such as accelerated speech rate or noise-masked speech (Hincapié-Casas et al., 2021; Nuttall et al., 2018). Furthermore, under challenging listening conditions, stronger right precentral activity has been directly associated with higher native consonant identification accuracy (Du et al., 2017; but see Du et al., 2014; Evans & Davis, 2015). Consistently, our univariate results highlighted the right precentral gyrus as a key region to correct categorization of native consonants as well as to non-native consonant perception under intact and noisy conditions. From a theoretical standpoint, the DIVA model of speech acquisition and production (Tourville & Guenther, 2011) posits a right-lateralized feedback control map in the premotor cortex, that transforms sensory error signals computed in bilateral auditory and somatosensory regions into corrective motor commands. These feedback-based corrections can be integrated into feedforward motor programs, supporting stable sensorimotor representations of speech sounds. Such representations may be engaged in the decoding of articulatory features during speech perception in both hemispheres, particularly under challenging listening conditions. The emphasis on the left motor cortex in the broader neuroimaging literature therefore likely stems from a methodological bias, as a substantial body of evidence, particularly from TMS studies, has restricted investigations to the left hemisphere (see Tseng et al., 2026 for a review).

### Limitations of the study

Our experimental stimulus set comprised consonants that naturally occur in French and Mandarin inventories but where the targeted phonetic features, in particular aspiration and voicing, were not represented equally. This potentially results in a partial confounding of segment identity and phonetic features. While an artificial, fully factorial design would allow us to isolate these dimensions completely, our methodological choice ensures that our findings reflect more ecologically valid speech processing. In natural language, features do not exist in isolation; rather, they are inherently tied to specific consonant classes.

While we should be cautious when assigning neural patterns to isolated phonetic features, our results offer a more realistic mapping of native and non-native speech sound processing in the brain. The use of a single speaker and vowel context, and of repeated consonant tokens, also limits the extent to which phonetic-feature representations can be dissociated from token-specific acoustic properties, despite the stimulus selection being guided by behavioral pilot studies and practical constraints of the fMRI protocol. The articulatory properties of the consonants used in the motor localizer correspond closely to the perceived labial and dental plosives but are less well matched to the fricative consonants. Future studies could consider alternative designs, such as using multiple speech targets (e.g., /b/, /d/, /n/, /l/) or non-speech articulator movements (e.g., Pulvermüller et al., 2006), to sample a broader range of articulatory properties and further disentangle the contributions of phonetic, articulatory, and perceptual factors to articulator-specific motor representations. Such designs could also incorporate behavioral tasks that independently assess consonant identification or phonetic-feature discrimination, allowing perceptual effects to be more clearly isolated from native and non-native language categorization processes and their associated decision-making demands. Finally, the interpretation of our RSA findings could be further strengthened by disentangling the potential contributions of acoustic and articulatory similarity to the neural representation of phonetic features. Future work may use more fully factorial stimulus sets with greater variability in speakers, vowels, and tokens. Complementary or causal methods in larger samples could furthermore help isolate these potential factors.

## CONCLUSION

The present fMRI study clarifies the role of the motor system in speech perception by mapping the transition from language-experience-dependent, articulator-specific motor representations to higher, language-independent phonetic feature representations. By combining complementary multivariate approaches, we demonstrate how the brain processes native and non-native speech sounds under varying listening conditions. To our knowledge, this study provides the first MVPA evidence that neural patterns during native speech perception map onto somatotopically-organized articulator-specific representations recruited during production. Crucially, this perception-production mapping is not invariant but constrained by the listener’s articulatory experience and selectively engaged under challenging conditions. Our RSA furthermore demonstrate that phonetic features across native and non-native speech sounds are robustly encoded within a sensorimotor network encompassing both auditory and motor regions. Altogether, these findings challenge the strict auditory and motor views of speech processing, supporting instead an integrated sensorimotor account. They suggest that the motor system is dynamically recruited to support speech perception in noise based on articulatory experience, consistent with a broader framework of embodied cognition, while also encoding stable higher-level phonetic features across languages.

## RESOURCE AVAILABILITY

### Lead contact

Requests for further information and resources should be directed to and will be fulfilled by the lead contact, Tzuyi Tseng.

### Materials availability

This study did not generate new unique reagents.

### Data and code availability

The original code and scripts are available on Gitlab at https://github.com/tzuyitseng/DecodPhone_fMRI and are publicly available as of the date of publication.

## ACKNOWLEDGMENTS

We thank F. Lamberton and D. Ibarrola for help with fMRI acquisitions and settings; the research assistants, J. Forano, A. Aymard and L. Demouliere for helping with the pilots and behavioral inclusion; G. Leroux for help with the server and *fMRIPrep* pipeline setup; B. Thirion, R. Gau, and H. Aggarwal for their help and suggestions regarding the Nilearn scripts; and R. Makine, S. Alouche, and J.-L. Borach for administrative support.

This project was supported by the French National Research Agency (ANR) for the AnchorFL project (ANR-19-CE28-0015; V.B., A.C.R. and C.B.), by the Appel à Projets Pluridisciplinaires Interne (APPI) from Université Lumière Lyon 2 (V.B., A.C.R. and C.B.), by Laboratory of Excellence ASLAN (ANR-10-LABX-0081) of Université de Lyon within the program “Investissements d’Avenir” (ANR-11-IDEX-0007; V.B., A.C.R., J.K., T.T.) of the French government operated by the ANR, and by the James S. McDonnell Foundation (JSMF; doi.org/10.37717/2021-3101; C.B.).

## Author contributions

T.T.: Methodology, Investigation, Data curation, Formal Analyses, Software, Visualization, Writing – Original Draft, Writing – Review and Editing. S.T.: Formal analyses, Software, Writing – Original Draft, Writing – Review and Editing. J.K.: Methodology, Formal analyses, Writing – Review and Editing. M.C.: Writing – Original Draft, Writing – Review and Editing. A.C.R., C.B. and V.B: Conceptualization, Methodology, Formal Analyses, Resources, Writing – Original Draft, Writing – Review and Editing, Supervision, Project Administration, Funding Acquisition.

## Declaration of interests

The authors declare no competing interests.

## STAR★Methods

### Experimental model and study participant details

A total of 42 healthy adult participants were recruited for the study, which consisted of a behavioral inclusion session and a fMRI session. Thirty-two participants passed the inclusion pre-test (i.e. ten participants did not pass the inclusion task, see Behavioral task screening and familiarization in Supplemental Information) and participated in the fMRI acquisition. Out of this sample, eight participants were excluded due to a withdrawal (n = 1), massive head motions (n = 1), and low behavioral performance during scanning (based on the same criterion used for inclusion, n = 6), resulting in 24 analyzed participants (17 females, age range 20 to 31 years old, age mean = 24.45 ± 3.12). All participants were right-handed (Edinburgh handedness inventory, Oldfield, 1971: 0.9 ± 0.12), French native monolingual speakers who had not acquired a second language before the age of 5. None had knowledge of or prior experience with Mandarin. An after-task language-background questionnaire indicated that, although some participants had incidental exposure to English or other languages (except Mandarin), none reported proficiency above the intermediate level (B1). Participants had normal pure-tone thresholds in both ears (-10 to 25 dB HL for 250 to 4,000 Hz, tested with the audiometer 600 M, Electronica Technologies). No known motor, linguistic, or neurological disorders were reported. All procedures were approved by an ethical committee (Comité de Protection des Personnes OUEST IV, 69HCL17_0359 NCT03223090). All participants signed a consent form before the experiment and were paid for their participation.

### Method details

#### Stimuli

Eight consonant-vowel (CV) syllables including four French tokens (/pa/, /ta/, /ʃa/ or /ʁa/) and their four Mandarin “counterparts” (/pʰa/, /tʰa/, /ʂa/ or /xa/) were recorded (16-bit quantization; 44.1 kHz sampling rate) in a sound-isolated booth by a female native bilingual French-Mandarin speaker. Mandarin and French, belonging to different language families, display both shared consonants and consonants that are language-specific. More specifically, these two languages allow to examine behavioral and cortical responses to consonants featuring either the same phonetic feature (e.g. for place of articulation: lips for the bilabials /p/ and /pʰ/, and tongue for the dentals /t/ and /tʰ/) or that differ on one or more features. Here, only a single phonetic feature differentiates the French and Mandarin consonants in three of the pairs (i.e. aspiration for the bilabial /p/-/pʰ/ and dental plosives /t/-/tʰ/, and tongue retroflection for the coronal fricatives /ʃ/-/ʂ/). By contrast, the dorsal fricatives (voiced uvular /ʁ/ and voiceless velar /x/) differ by two phonetic features, namely voicing and place of articulation (Figure S8, see measured acoustic properties in Table S3). To avoid potential confounds from vowel-related prosody or tone, we replaced the vowel /a/ from the originally-produced Mandarin tokens by the corresponding French /a/ in each consonant pair (/p/-/pʰ/, /t/-/tʰ/, /ʃ/-/ʂ/ and /ʁ/-/x/) using Praat (Boersma & Weenink, 2022). The stimulus set was selected based on several behavioral pilot studies that evaluated different combinations of ten consonants, six vowels, six speakers, and four signal-to-noise ratios (SNRs) (see Du et al., 2014; Osnes et al., 2011). These pilots guided the final selection of stimuli to maximize coverage of the relevant phonetic-feature dimensions across the two languages while preserving the phonological structure of each language to maintain ecological validity. Consequently, some feature contrasts (e.g., aspiration and voicing) are language-specific and occur only within particular consonant classes, precluding a fully factorial manipulation of all phonetic dimensions. Certain phonetic features are therefore unevenly distributed across the stimulus set, with aspiration primarily characterizing plosives and voicing tied to specific fricatives. Each syllable lasted approximately 500 ms and was fixed at 80 dB in terms of average root mean square sound pressure level (SPL). To create the noisy versions of the stimuli, each of the eight syllables was masked by a 500 ms pink noise segment at 72 dB SPL (i.e. SNR = 8, see Du et al., 2014) using a Praat script (Petersen, 2004). We then generated triplets of syllables as stimuli of interest: each of the intact (non-noisy) and noisy syllables were reduplicated three times and concatenated every 500 ms as a triplet. The short interstimulus interval (ISI; time interval between the offset of a syllable and the onset of the next one) in the triplet was filled either with silence or pink noise (for the noisy version). In total, the experiment included 16 different stimuli (8 syllable triplets × 2 noise conditions, intact and noisy respectively) that were presented at 80 dB SPL.

#### Task and experimental procedure

Participants who passed the behavioral inclusion test (see *Behavioral task screening and familiarization* in *SI*) underwent the fMRI session on a different day. The task in the scanner was a two-alternative forced choice (2AFC) task in which participants were required to listen to triplets of CV syllables (e.g., /pa pa pa/) in the two languages and categorize the heard consonant as native or non-native. It consisted of 6 experimental blocks, each including 48 trials (8 triplets of syllables × 2 noise conditions × 3 repetitions), resulting in a total of 288 trials for a duration of one hour (Figure S9). Each block began with a 48-s period with a grey fixation point displayed at the center of the screen, used as a baseline. This long baseline duration is related to the sparse sampling fMRI design (see *MRI Acquisition*). Each trial then started with a white fixation point at the same location, shown for a duration jittered between 500 and 750 ms, which was followed by the auditory stimulus (i.e. a 1500 ms triplet of CV syllables) and a 1500 ms window for response. Participants responded by pressing one of two buttons using their left index or middle finger, corresponding to icons on the screen: a French flag for native consonants and an earth icon for non-native consonants. The association between the button press (finger used) and the icons on the screen was counterbalanced across participants. These icons stayed on the screen until the participants’ response or for a maximum of 1500 ms in case of no response, and they were replaced by a grey fixation point (4500 ms) before the next trial.

Given the use of a sparse sequence for MRI, featuring interspersed silent and acquisition periods (see MRI Acquisition), each trial was structured as follows: a jittered fixation period (500/750 ms), auditory stimulus presentation (1500 ms), and a behavioral response window (1500 ms). This silent period was immediately followed by the fMRI acquisition period (4500 ms), resulting in a total trial duration (corresponding to intertrial interval here) varying between 8000 or 8250 ms (Figure 7). Auditory stimuli were presented through insert earphones (Sensimetrics Model S14 with standard Comply Foam Canal Tips) to reduce the impact of the scanner noise on the perceptual categorization task.

The consonant categorization task (2AFC) was followed by a motor localizer task, in which the participants were required to perform small movements of the lips (silently articulating “ma”), tongue (silently articulating “la”), as well as left and right index fingers (tapping movements) using continuous MRI sampling. The purpose of the motor localizer was to isolate independent, articulator-specific motor representations for the lips and the tongue. We therefore used these French native consonants to cue the clear somatotopic markers for lip (/m/) and tongue (/l/) movements, respectively, while minimizing laryngeal confounds during production. There was one block, which included 6 motor trials for each body part, resulting in a total of 24 trials randomized across participants. Each trial began with a 15000 ms baseline recording (with a grey fixation point displayed on the screen), followed by 2000 ms of written instructions, 10000 ms of movements with a white fixation point, and 6000 ms of rest with an orange fixation point to let the BOLD signal come back to baseline levels. The fMRI session ended with a 5-minutes T1-weighted anatomical acquisition, for a total duration of 1 hour and 30 minutes for each participant.

### Quantification and statistical analysis

#### Behavioral Data Acquisition & Analysis

The behavioral task was programmed and participants’ responses were recorded with OpenSesame 3.3.11 (Mathôt et al., 2012; Mathôt & March, 2022). Behavioral accuracy (% of correct consonant categorization) in the 2AFC task was assessed using a repeated measures ANOVA including language (native vs. non-native), noise (intact vs. noisy) and consonant (bilabials /p-pʰ/, dentals /t-tʰ/, coronal fricatives /ʃ-ʂ/ vs. dorsal fricatives /ʁ-x/) as within-subject factors, and subjects as a random factor. Post-hoc tests (Tukey’s HSD) were conducted to assess significant interactions between the factors. Statistical analyses were run in Python 3.11.5.

#### MRI Acquisition

Participants’ brain activity was recorded with a Siemens Prisma 3T scanner (Siemens Medical Systems, Erlangen, Germany) with a 64-channel head/neck receive-array coil, using a multiband gradient echo planar imaging (EPI) sequence. Functional T2*-weighted images were acquired applying a sparse imaging sequence (Hall et al., 1999; Peelle et al., 2010), where speech stimuli are presented during silent TRs where no functional images are acquired so as to reduce scanner noise (Pulvermüller et al., 2006; Schomers & Pulvermüller, 2016). Brain activity to these stimuli is then captured with subsequent acquisition TRs following the timing of the hemodynamic response. Here, each sequence included three silent TRs, interspersed with three acquisition TRs (TR = 1500 ms). Six runs of sparse sampling were used for the consonant categorization task (2AFC). One run of continuous sampling, consisting exclusively of acquisition TRs, was applied for the motor localizer task. The same parameters were set for both sparse and continuous functional acquisition runs (echo time (TE) = 30 ms; TR = 1500 ms; flip angle = 75°; slice thickness = 2.7 mm; voxel size = 2.69 × 2.69 × 2.7 mm). Anatomical T1-weighted images were acquired with a 1-mm isotropic voxel and a generalized auto-calibrating partially parallel acquisition (GRAPPA) (TE = 2.8 ms, TR = 2400 ms). Overall, 158 volumes were acquired per run in the consonant categorization task (for a total of 948 volumes over the six runs) and 300 volumes were acquired in the motor localizer task.

#### fMRI Preprocessing

Preprocessing of the fMRI data was performed using the *fMRIPrep* pipeline version 23.0.1 (RRID:SCR_016216; Esteban et al., 2019). Functional data were motion corrected by volume-realignment using MCFLIRT (FSL 6.0.5.1:57b01774; Jenkinson et al., 2002) and registered to the MNI152NLin2009cAsym standard space template. BOLD runs were slice-time corrected to 0.699 s (the actual slice acquisition ranging from 0 to 1.4 s) using 3dTshift from AFNI (RRID:SCR_005927; Cox & Hyde, 1997).

#### Univariate Analyses

The univariate analyses were conducted using the Nilearn 0.10.2 library (Abraham et al., 2014) in Python. Six head motion parameters (3 translation and 3 rotation parameters) and a nuisance denoising regressor (Caballero-Gaudes & Reynolds, 2017) were included in a general linear model (GLM) as regressors of no interest. This regressor included the additional 3 temporal derivatives of each head motion parameter, 5 white matter principal components (PCs), and 5 cerebrospinal fluid PCs, which were available for all participants across runs. The hemodynamic response function (HRF) was modelled using the canonical Statistical Parametric Mapping (SPM) HRF (a double-gamma function, Friston et al., 1998). A high-pass filter of 128 s allowed to remove low-frequency signal drifts and the time of the reference slice was adjusted based on the parameter used in the slice timing preprocessing. The imaging data were smoothed with a 6-mm full width at half maximum (FWHM) kernel. For the consonant categorization data, a modulation regressor for behavioral accuracy was applied to include only trials with correct responses, excluding those with incorrect responses (see Figure S4 for Brain activations computed without applying the modulation regressor). Considering the ∼4 to 6 s delay of the BOLD signal from stimulus onset (Perrachione & Ghosh, 2013), only the functional data from the last two acquisition TRs after stimulus presentation (i.e. data included from 6 s after the fixation onset, with fixation jittered between 0.5 to 0.75 s) were included for further analyses. The data were then fitted into the first-level GLM to compute the contrasts of the 8 consonants (4 tokens by language) and 2 noise conditions.

To assess the overall activation for processing consonants in each language and each noise condition, contrasts against the baseline were calculated (Language and Noise Contrasts) as follows:

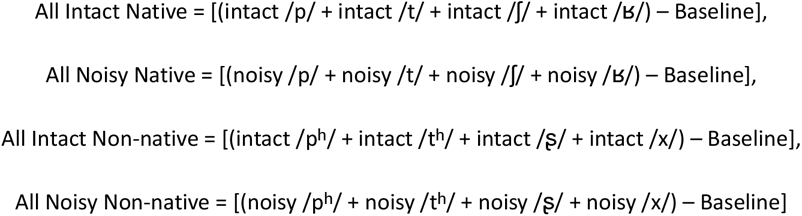

We also evaluated the main effects of language and noise independently with the following contrasts (Main Effect Contrasts):

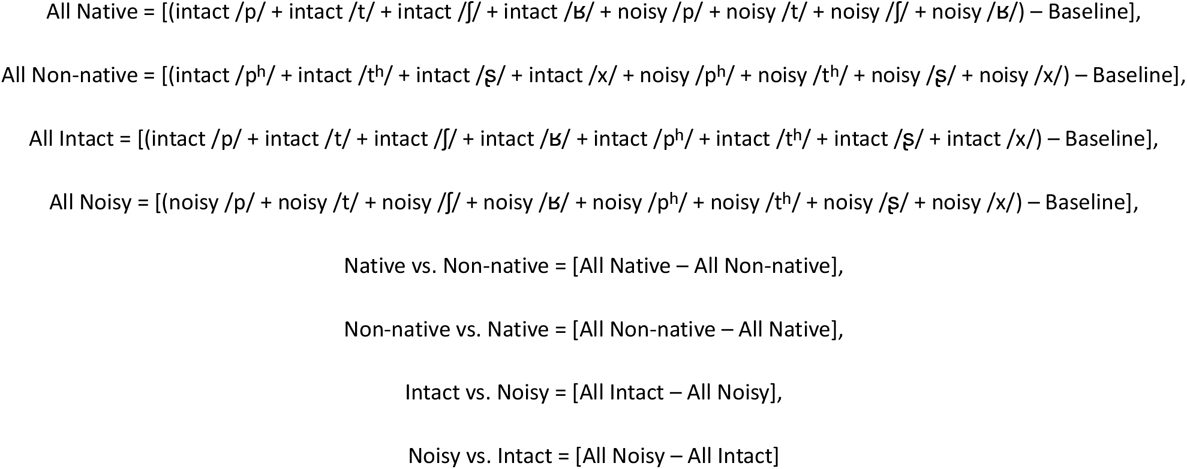

For the group-level analysis, we conducted a within-subject paired two-sample t-test for each of the contrasts computed at the first level (see Language and Noise Contrasts and Main Effect Contrasts above). To further explore the relationship between brain activation and behavioral performance, we additionally applied individual mean accuracy for consonant categorization (across trials) as a regressor of interest for the Main Effect Contrasts (All Native, All Non-native, All Intact and All Noisy) in the paired two-sample t-test.

Statistical inferences for the group-averaged activation were performed using a map-wise *p*-value threshold corrected for multiple comparisons with the False Discovery Rate (FDR, see Table S1, Figure S1, S2 and S3 for the univariate results).

### Multivariate Pattern Analysis

#### Cross-Modal Classification

To examine the motor somatotopy of consonant perception, we performed a cross-modal classification analysis to test for shared neural patterns in the motor cortex between articulatory lip and tongue movements and perception of bilabial and dental consonants. More specifically, we assessed whether a classifier trained on the motor neural patterns could predict the speech perception patterns.

The analysis used t-maps derived from motor localizer trials (lip and tongue movements) and consonant perception blocks (bilabial and dental consonants). The preprocessing steps for the cross-modal decoding for each participant were similar to the univariate first-level analysis but without applying any smoothing nor the behavioral modulation regressor. For the motor localizer data, all six trials of the two body part movements were fitted into the first-level GLM to compute the contrast activations of each articulator of interest (lips or tongue) against the baseline to localize the corresponding articulatory regions, yielding 6 t-maps for each contrast per participant:

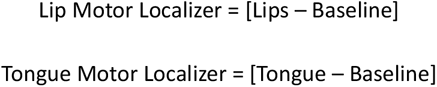

Similarly, for the speech perception data, contrasts of activation for consonant perception against the baseline (2 noise x 2 languages x 4 consonant conditions) were computed from each block (n = 6), yielding 6 t-maps for each contrast per participant (Consonant Perception Contrasts):

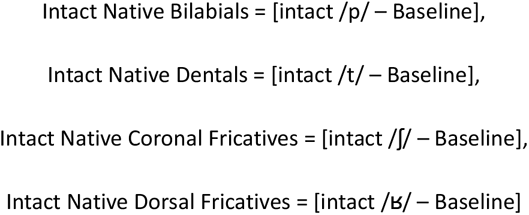

The same rationale was applied to compute contrasts for the Noisy Native, Intact Non-native and Noisy Non-native conditions (Table S4). Each of these consonant perception t-maps contained the averaged activations from the 3 repeated trials for each condition. Only bilabial and dental consonant perception data were used for classification. This led to 12 t-maps (6 runs x 2 consonants) for each language and perceptual condition (intact or noisy; e.g., 6 intact native bilabial perception t-maps, 6 noisy non-native dental perception t-maps, etc.).

The classification was conducted within individual spherical regions of interest (ROIs), using a linear support vector machine (SVM) classifier implemented in Nilearn. Each individual spherical ROI in each hemisphere was defined as a 10 mm sphere (mean voxel numbers = 190) around the peak coordinate (Table S5 and Figure S10) of the Mouth Motor Localizer contrast. This contrast was computed to isolate motor regions recruited for movements of both articulators of interest (lips and tongue), distinct from regions associated with finger movements, so to specifically identify articulator-specific representations in the motor cortex:

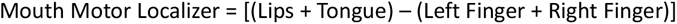

Before decoding, to minimize the univariate differences between the two modalities (articulatory movements and speech perception) and the two scan sequences (continuous and sparse sampling), we normalized the data (demeaning and z-scored standardization) of each t-map in each modality for each participant (Rezk et al., 2020). A within-modal classifier was first trained and tested on the motor data to discriminate the neural patterns of lip and tongue articulatory movements. A leave-one-trial-out cross-validation scheme was implemented, which led to 6 folds of cross-validation. In each fold, we trained the classifier on N-1 trials (10 t-maps for both classes, lips and tongue, 5 trials for each class) and tested it on the two t-maps from the two left-out trials (one for each class). The classifiers with above-chance within-modal decoding (chance level = 0.5) were used for cross-modal decoding, namely we additionally tested them on the activations elicited by the perception of bilabial and dental consonants in the 2AFC task (6 runs). The perception data for each language and noise condition were tested by the same classifier separately, i.e. each cross-modal decoding analysis was implemented on one of the four speech conditions (intact native, noisy native, intact non-native and noisy non-native) with 12 t-maps (6 t-maps x 2 classes: bilabials vs. dentals). An additional cross-modal decoding was similarly conducted, as a control, using the same classifier to test on the activations elicited by the perceived fricatives (12 t-maps: 6 t-maps x 2 classes: coronal vs. dorsal fricatives; see Figure S5). We obtained a classification accuracy score for each participant by averaging the classification accuracies of all cross-validation folds for each condition of each modality (i.e. movement and perception) and hemisphere. We then calculated group mean accuracy for each condition in each modality and hemisphere. Outliers (±3 SD from the mean, n = 4) were identified based on z-scores across participants and excluded prior to statistical analyses. Statistical significance was assessed by one-sample permutation tests against chance level with FDR correction for multiple testing correction across the four speech conditions in the two hemispheres.

### Representational Similarity Analysis

Through RSA (Kriegeskorte et al., 2008), we tested the similarity between the neural patterns induced by the different conditions, on the basis of a priori theoretical models. We performed a searchlight RSA to assess whether the examined representational similarity structure of phonetic features was encoded in predefined ROIs during consonant perception. The theoretical models were conceptualized by modeling the similarity between the phonetic features of the heard consonants across languages and perceptual conditions. The theoretical representational dissimilarity matrices (RDMs; Figure 3) were expressed as a set of 16 x 16 matrices corresponding to the 16 experimental conditions (8 consonants x 2 noise conditions). Each cell in the matrix represents the predicted similarity (converted as dissimilarity for computations) between the activity patterns of two conditions. A value of +1 (blue tiles) indicates increased dissimilarity (i.e. decreased similarity) between conditions, while -1 values (red tiles) indicate decreased dissimilarity (i.e. increased similarity). Elements of the matrices that were irrelevant to the tested hypotheses were excluded (N/A values, green tiles). For example, diagonal values were excluded to avoid high correlation between the exact same conditions (Evans & Davis, 2015; Ritchie et al., 2017). We designed four theoretical RDMs to assess the neural representation of four phonetic features: place of articulation (Figure 3A), manner of articulation (Figure 3B), aspiration (Figure 3C) and voicing (Figure 3D). Consonants with different phonetic features in each model were considered to be represented by dissimilar neural patterns (+1 in the model), irrespective of language and noise (e.g., native bilabial /p/ and non-native dental /tʰ/ for place; native plosive /p/ and native fricative /ʁ/ for manner; native unaspirated /p/ and non-native aspirated /tʰ/ for aspiration; native voiceless /ʃ/ and native voiced /ʁ/ for voicing). On the contrary, consonants with the same phonetic features of interest were given -1 value for similarity of their neural patterns (e.g., native bilabial /p/ and non-native bilabial /pʰ/ for place; native fricative /ʃ/ and non-native fricative /x/ for manner; native unaspirated /p/ and native unaspirated /t/ for aspiration; native voiceless /ʃ/ and non-native voiceless /x/ for voicing). For place and manner of articulation, we took all the consonants (8 consonants x 2 noise conditions) into account. For aspiration, we modelled only the four plosives (/p/, /pʰ/, /t/ and /tʰ/), while for voicing, we modelled only the four fricatives (/ʃ/, /ʂ/, /ʁ/ and /x/) to reduce comparisons across unbalanced conditions. An additional theoretical RDM based on language identity was computed (Figure S7A) to test whether neural representations of speech sounds are organized by language boundaries. Consonants belonging to the same language were assigned a value of -1 for similarity of their neural patterns (e.g., native /p/ and native /ʃ/), whereas consonants belonging to different languages were considered to be represented by dissimilar neural patterns (e.g., native /t/ and non-native /tʰ/).

For the speech perception data used in RSA, the preprocessing steps were similar to those in the cross-modal decoding analysis but included all consonants. A single t-map for each of the Consonant Perception Contrasts (see Cross-modal Classification) was computed across the 6 runs. We defined seven bilateral ROIs based on previous neuroimaging findings on speech processing (Archila-Meléndez et al., 2018; Arsenault & Buchsbaum, 2015; Correia et al., 2015; Du et al., 2014; Evans & Davis, 2015) using the AAL atlas (Tzourio-Mazoyer et al., 2002). These ROIs included the bilateral precentral and postcentral gyri, inferior frontal gyrus (IFG), insula, supramarginal gyrus (SMG), and superior and middle temporal gyri (STG and MTG).

The 7 ROIs were combined into a single ROI for the searchlight approach (see Correia et al., 2015). A 10 mm spherical radius moved across each voxel in the combined ROI, with the voxel serving as the center of the searchlight sphere (n = 14,199 voxels). At each searchlight center, a neural RDM was generated and correlated to a theoretical RDM using an open-source Python toolbox (rsatoolbox.readthedocs.io, Nili et al., 2014). The searchlight was performed across the entire combined ROI mask, meaning that searchlights could cross the boundaries between the 7 ROIs included in this mask. Specifically, each neural RDM was calculated from the pairwise correlation between neural patterns of each condition (n = 16) using cosine distance, resulting in a 16 x 16 pairwise matrix. We then calculated the correlation between each of the theoretical RDMs (predicted similarity) and the neural RDMs (observed similarity) using Spearman’s rho (ρ). The better the theoretical models fit the empirical data (higher rho values), the stronger the evidence that the modelled phonetic features are encoded in the observed neural activity. We constructed an individual brain map by projecting the Spearman’s rho values of each voxel back onto each subject’s anatomical space. To perform a group analysis, we then entered these individual brain maps into a second-level model. The voxel-level significance was computed with a non-parametric permutation test (n = 10,000). The corresponding voxel-wise p-values were corrected using an FDR threshold at 0.05. Such correction was applied separately for each of the four models. We further conducted a repeated-measures ANOVA to investigate the specificity of phonetic feature encoding across STG and precentral gyri (Evans & Davis, 2015) and hemispheres. Spearman’s rho values extracted from these two regions were included in the model, with ROI (STG vs. precentral gyri), Hemisphere (left vs. right) and Phonetic feature (place of articulation, manner of articulation, aspiration vs. voicing) as fixed factors, and subjects as a random factor. This analysis was conducted for phonetic features meeting a predefined criterion from the preceding RSA (i.e. features represented in both hemispheres and across both ROIs). Post-hoc paired t-tests were conducted to assess significant interactions.

### Key resources table

| REAGENT or RESOURCE | SOURCE | IDENTIFIER |
|---|---|---|
| **Deposited data and scripts** |  |  |
| *fMRIprep* QC report | This paper | github.com/tzuyitseng/DecodPhone_fMRI/fMRIprep_reports |
| fMRI first-level brain maps | This paper | github.com/tzuyitseng/DecodPhone_fMRI/first_level |
| fMRI scripts | This paper | github.com/tzuyitseng/DecodPhone_fMRI/fMRI_scripts |
| **Software and algorithms** |  |  |
| Praat | Boersma & Weenink, 2022 | www.fon.hum.uva.nl/praat/ |
| Opensesame | Mathôt et al., 2012;<br>Mathôt & March, 2022 | github.com/psychopy/psychopy |
| fMRIPrep 23.0.1 | Esteban et al., 2019 | fmriprep.org/en/stable/ |
| Nilearn 0.10.2 | Abraham et al., 2014 | nilearn.github.io |

## Supplemental information

Document S1. Supplemental methods, Figures S1–S10, and Tables S1–S5

## Notes

https://github.com/tzuyitseng/DecodPhone_fMRI

### Competing Interest Statement

The authors have declared no competing interest.

### Summary of Updates

This revised version of the manuscript has been prepared in response to the Editorial Decision on Manuscript ISCIENCE-D-26-07134. The revisions have been carefully implemented to enhance the clarity, accuracy, and overall quality of the manuscript.

