## Supplemental information for "Decoding Articulatory and Phonetic Features: Somatotopic and Sensorimotor Representations in Native and Non-native Consonant Perception"

### RESULTS

#### fMRI Univariate activations

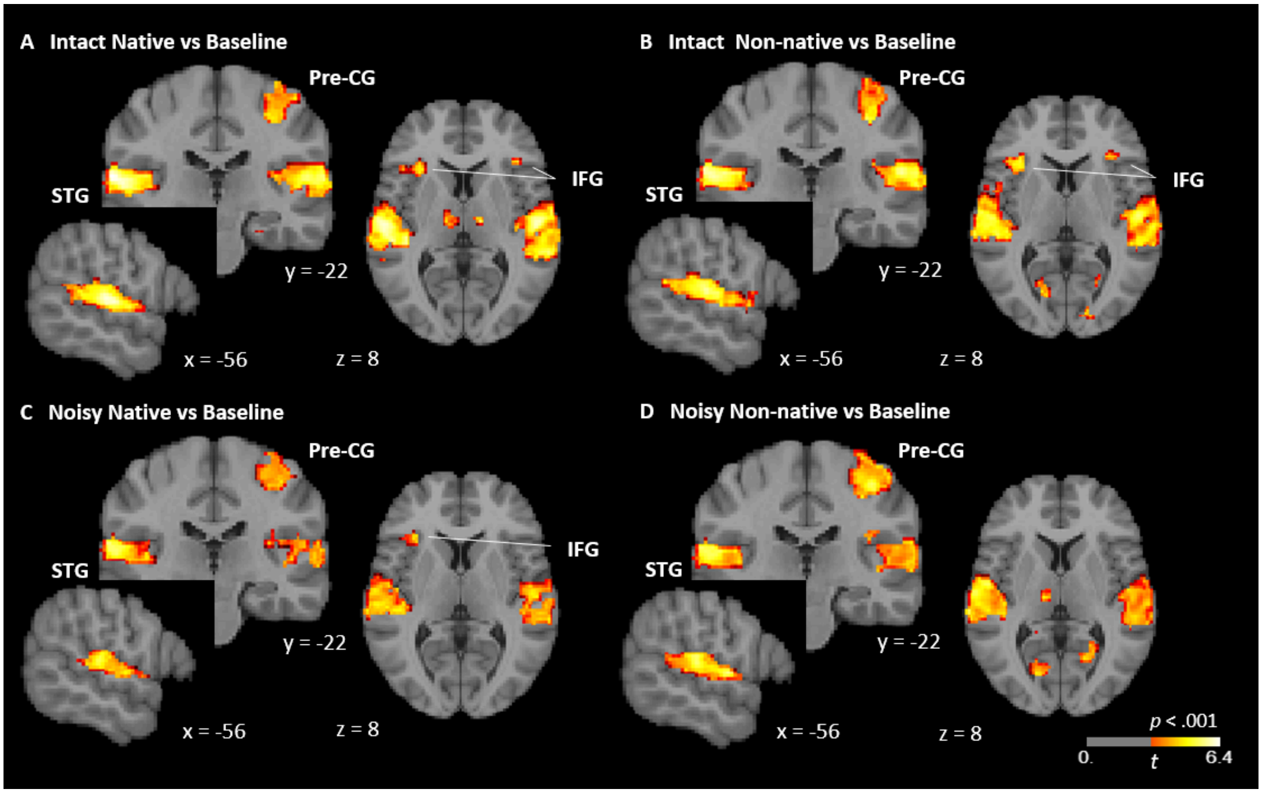


**Figure S1.** Brain activations for consonant perception in the native or non-native language and under intact or noisy conditions. Brain maps show the brain activation (against baseline) induced during the two-alternative forced choice (2AFC) categorization task for (A) intact native vs. baseline, (B) intact non-native vs. baseline, (C) noisy native vs. baseline, and (D) noisy non-native vs. baseline (see Table S4 for cluster peak coordinates). The perception of consonants to be categorized elicited activations in the bilateral STG and STS and in the right precentral gyrus in all conditions when compared to baseline. Examining the intact conditions only in (A) and (B), we observed that both native and non-native consonants induced additional bilateral activations in the IFG compared to baseline. When noise was added in (C) and (D), IFG activation was restricted to the left hemisphere for native consonants, whereas no significant activation was observed in this region for non-native consonants. Activation thresholds were set at $p < .001$, FDR-corrected, with a minimum cluster size of 195 $mm^3$. The color bar represents t-values exceeding the activation threshold. Pre-CG: precentral gyrus. STG: superior temporal gyrus. STS: superior temporal sulcus. IFG: inferior frontal gyrus.

**Table S1.** Cluster peak MNI coordinates of brain activation for consonant perception in the native or non-native language and under intact or noisy conditions. Pre-CG: precentral gyrus. Post-CG: postcentral gyrus. STG: superior temporal gyrus. IFG: interior frontal gyrus. INS: insula.

| Region | X | Y | Z | Peak Stat | Cluster Size ($mm^3$) |
|---|---|---|---|---|---|
| **Intact Native vs Baseline** | | | | | |
| STG | -64 | -25 | 11 | 6.38 | 15164 |
| | 54 | -17 | 5 | 6.04 | 20153 |
| IFG/INS | -37 | 21 | -6 | 4.84 | 2015 |
| | 35 | 24 | -3 | 4.34 | 1819 |
| Pre-CG | 35 | -25 | 48 | 4.69 | 4676 |
| **Intact Non-native vs Baseline** | | | | | |
| STG | -64 | -22 | 13 | 6.07 | 15535 |
| | 57 | -22 | 13 | 5.84 | 18196 |
| Pre-CG | 35 | -22 | 46 | 5.01 | 5126 |
| IFG/INS | -35 | 24 | 5 | 4.98 | 1193 |
| | 35 | 26 | 8 | 4.24 | 665 |
| **Noisy Native vs Baseline** | | | | | |
| STG | -64 | -22 | 13 | 5.92 | 10448 |
| | 62 | -9 | 0 | 4.95 | 11035 |
| Pre-CG | 33 | -30 | 59 | 4.65 | 5830 |
| IFG/INS | -35 | 24 | 5 | 4.11 | 547 |
| **Noisy Non-native vs Baseline** | | | | | |
| STG | -59 | -6 | 3 | 5.56 | 12953 |
| | 65 | -9 | 0 | 5.00 | 14792 |
| Pre-/Post-CG | 41 | -25 | 54 | 5.42 | 9900 |

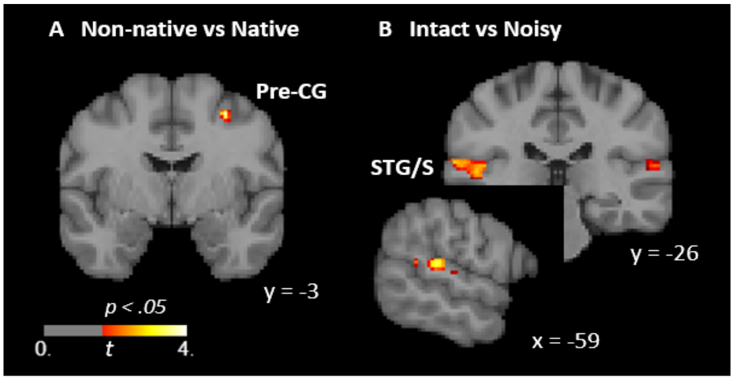


**Figure S2.** Brain activations elicited by the categorization of non-native or intact consonants. Brain maps show the significant clusters identified from the (A) Non-native vs. Native contrast (see *Methods*) comparing intact and noisy non-native against intact and noisy native consonants in the right precentral gyrus (pre-CG; peak MNI at 33, -3, 51), (B) Intact vs. Noisy contrast comparing intact native and non-native against noisy native and non-native consonants in the bilateral superior temporal gyrus/sulcus (STG/STS; peak MNI at, respectively, -59, -33, 13 and 57, -38, 16). No significant activation was observed for the reverse contrasts. The bilateral STG was more engaged when participants categorized consonants under easier (intact) perceptual conditions, whereas the right pre-CG was more activated for the non-native language. For display purposes, brain maps are set to an FDR-corrected threshold p < .05 and a cluster size > 195 $mm^3$. The color bar indicates the activation t-values above threshold.

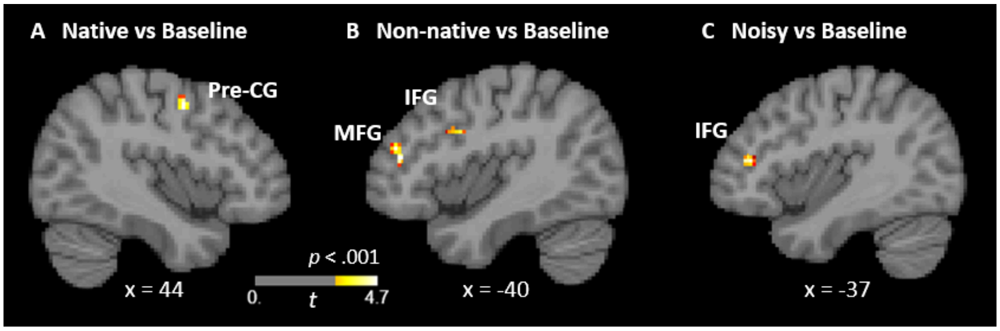


**Figure S3.** Cortical regions showing significant correlation between behavioral accuracy in the 2AFC categorization task and BOLD signal when contrasting consonant perception with baseline. Brain maps show the clusters identified from the (A) All Native contrast (see Methods section) comparing intact and noisy native consonants against baseline in the right precentral gyrus (pre-CG; peak MNI at 43, -3, 48). The better participants were at categorizing consonants in their native language, the more this right precentral region was engaged; (B) All Non-native contrast comparing intact and noisy non-native consonants against baseline in the left inferior and middle frontal gyri (IFG and MFG; peak MNI at, respectively, -48, 10, 30 and -42, 42, 13); (C) All Noisy contrast comparing noisy native and non-native consonants against baseline in the left IFG (peak MNI at -37, 39, 13). This indicates that the left IFG was more strongly activated for better behavioral performance under the noisy perceptual condition. No significant cluster of activation correlated with behavioral accuracy for the All Intact contrast comparing native and non-native consonants with baseline. Brain maps are restricted with the threshold at FDR-corrected $p < .001$ and cluster size > 195 $mm^3$. The color bar indicates the activation t-values above threshold.

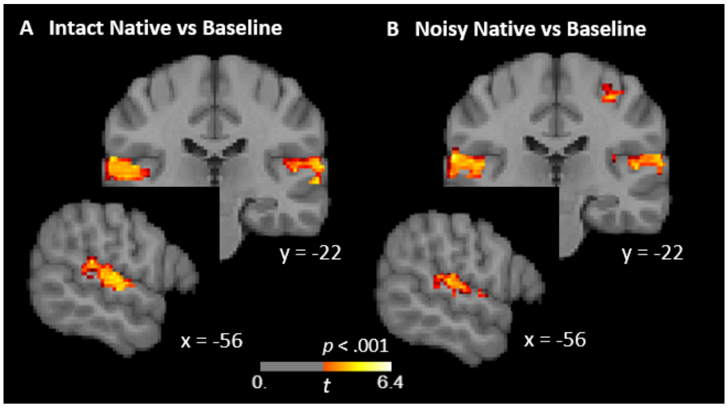


**Figure S4.** Brain activations during the 2AFC consonant categorization task considering both correct and incorrect responses. These maps were computed without applying the modulation regressor aimed at excluding trials with incorrect responses (as in Figure S1). Activity in the right precentral gyrus was no longer observed for intact native consonants (A; All Intact Native contrast in the Methods section) and was reduced for noisy native consonants (B; All Noisy Native contrast) compared to when the modulation regressor was applied (Figure S1). This suggests that right precentral activity was associated with correct native consonant categorization, in line with other studies (Callan et al., 2010; Smalle et al., 2015).

**fMRI MVPA – Cross-modal decoding**

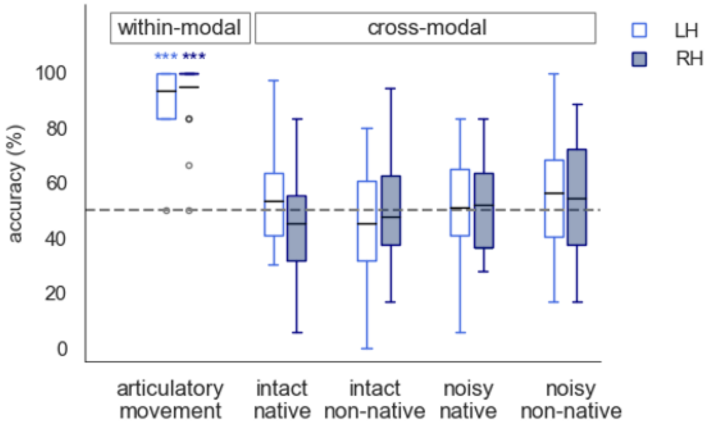


**Figure S5.** MVPA classification accuracy of motor patterns (within modality, first two boxplots on the left) for lip and tongue articulatory movements, and cross-modal pattern classification (the rest 8 boxplots on the right) for the perception of the native and non-native coronal (/ʃ/, /ʂ/) and dorsal (/x/, /ʁ/) fricatives in the intact and noisy conditions. For each condition, the first boxplot shows the mean classifier accuracy in the precentral ROI of the left hemisphere (LH, light blue unfilled boxes); the second one shows the accuracy in its counterpart in the right hemisphere (RH, dark blue filled boxes). The black horizontal line in each box indicates the group mean accuracy for the corresponding condition. Asterisks (***p < .001) indicate statistical significance in the one-sample t-test against chance level (50%) with FDR correction and multiple testing correction for the conditions in the two hemispheres. No significant classification accuracy against chance level was observed for any of these consonant perception conditions.

#### fMRI MVPA – RSA

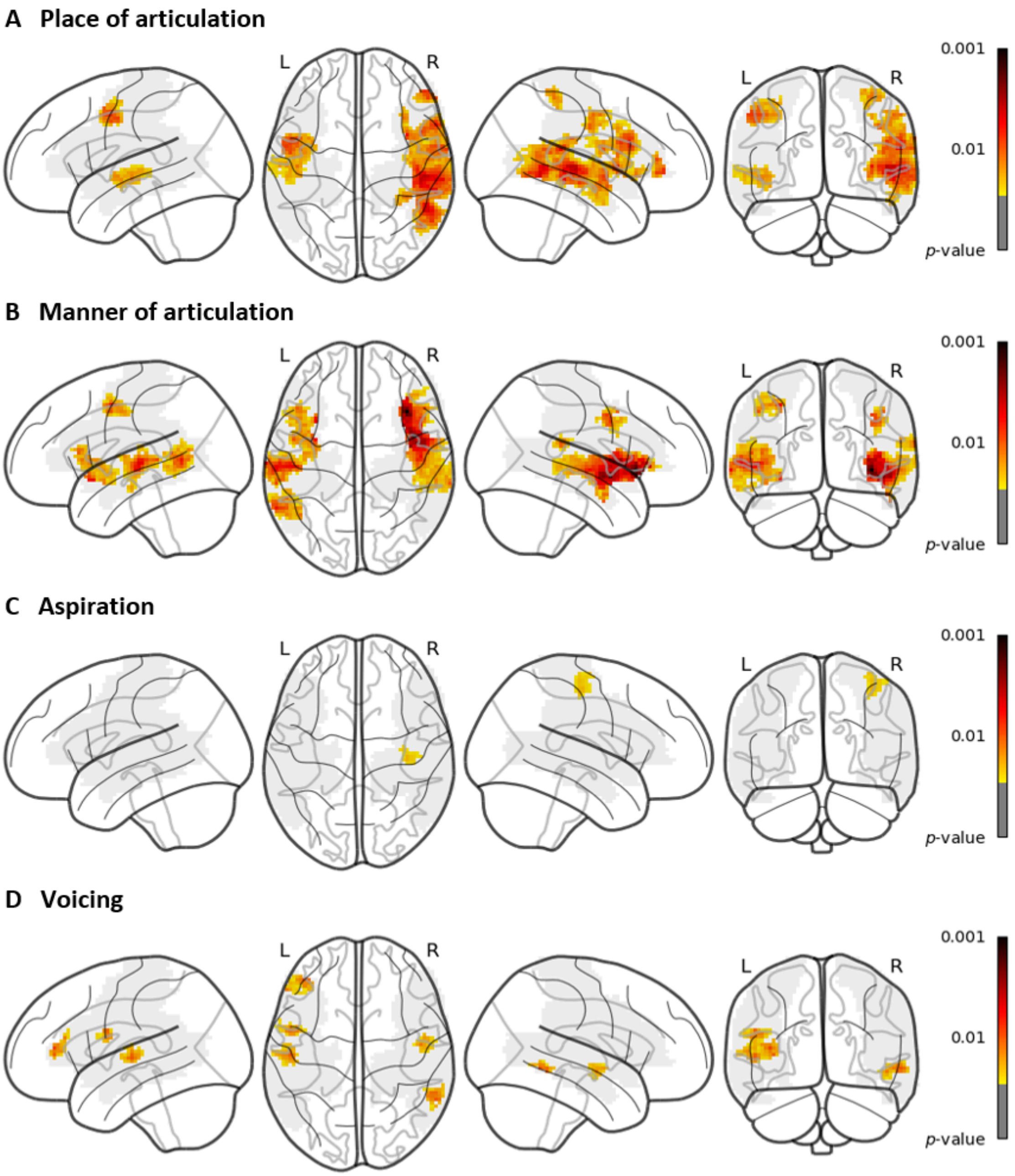


**Figure S6.** Searchlight RSA brain maps illustrating the similarity in neural activity patterns for consonants sharing the same phonetic feature during perception: (A) place of articulation, (B) manner of articulation (C) aspiration, and (D) voicing. Brain maps show the significant clusters after permutation testing thresholded at $p < .05$ (FDR-corrected) with a minimum cluster size of 780 $mm^3$. Shaded grey areas indicate the regions of interest. The color bar indicates the activation t-values above threshold ($p < .05$) in each condition.

**Table S2**. Peak MNI coordinates of neural activity patterns found in searchlight RSA brain maps. IFG: interior frontal gyrus. INS: insula. Pre-CG: precentral gyrus. Post-CG: postcentral gyrus. MTG: middle temporal gyrus. SMG: supramarginal gyrus. STG: superior temporal gyrus.

| Region | X | Y | Z | Peak Stat | Cluster Size ($mm^3$) |
|---|---|---|---|---|---|
| **Place of articulation** | | | | | |
| Pre-CG | -51 | 2 | 46 | 3.32 | 4206 |
| | 43 | -14 | 38 | 2.84 | 2054 |
| Post-CG | -51 | -6 | 40 | 2.89 | 430 |
| | 68 | -3 | 19 | 3.00 | 1369 |
| IFG | 60 | 16 | 24 | 3.71 | 4695 |
| INS | -43 | -3 | -3 | 2.43 | 450 |
| | 38 | 6 | 5 | 3.58 | 2837 |
| SMG | 70 | -28 | 27 | 2.98 | 626 |
| STG | -45 | -17 | -6 | 2.72 | 2563 |
| | 52 | -25 | 3 | 4.28 | 15868 |
| MTG | -59 | -17 | 0 | 2.29 | 1467 |
| | 52 | -52 | 0 | 4.03 | 12816 |
| **Manner of articulation** | | | | | |
| Pre-CG | -32 | 0 | 48 | 3.84 | 2406 |
| | 41 | 2 | 35 | 3.31 | 1193 |
| IFG | -45 | 24 | 11 | 2.88 | 1976 |
| | 38 | 32 | 0 | 4.03 | 2915 |
| INS | -32 | 13 | -11 | 3.76 | 1917 |
| | 38 | 21 | -3 | 6.13 | 7200 |
| STG | -48 | -22 | 3 | 3.90 | 2582 |
| | 46 | -3 | -8 | 4.88 | 8804 |
| MTG | -51 | -22 | 0 | 3.75 | 3913 |
| | 49 | -28 | -6 | 2.36 | 1310 |
| **Aspiration** | | | | | |
| Pre-CG | 38 | -19 | 57 | 2.10 | 1252 |
| **Voicing** | | | | | |
| Pre-CG | -51 | 2 | 24 | 2.80 | 841 |
| IFG | -40 | 42 | 11 | 3.11 | 1506 |
| STG | -51 | -17 | 3 | 2.91 | 1584 |
| | 49 | -9 | -8 | 2.61 | 1076 |
| MTG | 60 | -52 | -6 | 2.93 | 802 |

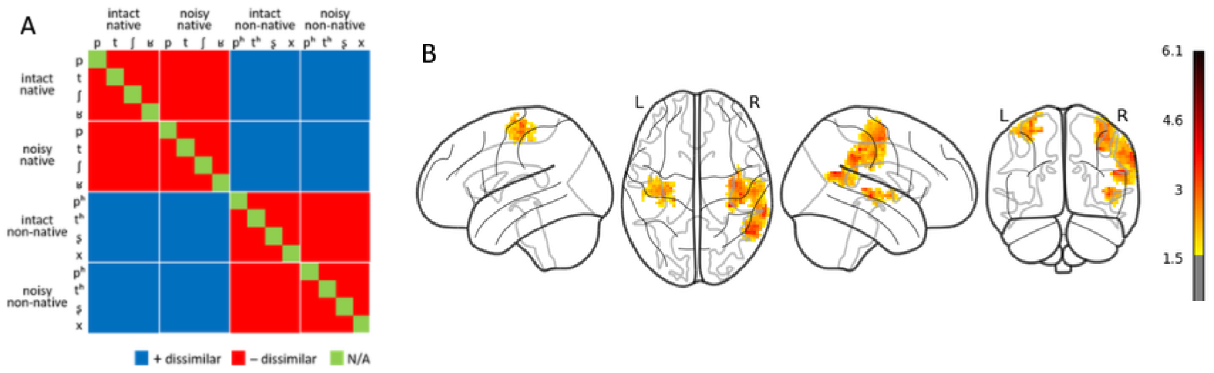


**Figure S7. RSA based on language identity.** (A) Language representational dissimilarity matrix (RDM). Theoretical model RDM predicting the similarity structure of the neural patterns based on native and non-native language. Each tile represents the hypothesized similarity/dissimilarity between the neural patterns for two consonants, in the intact native, noisy native, intact non-native, noisy non-native conditions. Pairs of consonants from the same language were assigned -1 value (similar neural patterns, red tiles), whereas those from different languages were assigned +1 values (dissimilar patterns, blue tiles). Diagonal values (N/A values, green tiles) were excluded to avoid high correlation between the exact same conditions (Evans & Davis, 2015). (B) Searchlight RSA brain maps associated with language identity. Searchlight RSA brain maps show the involvement of a network distributed over bilateral pre- and post-central gyri, as well as the right supramarginal, superior and middle temporal gyri. Neural patterns of these identified clusters are sensitive to perceived consonants that reflect within-language similarity and between-language dissimilarity. Brain maps show the significant clusters after permutation test with FDR-corrected $p < .05$ and a cluster size > 780 $mm^3$. The color bar indicates the activation *t*-values above the threshold.

### METHODS

#### Stimuli

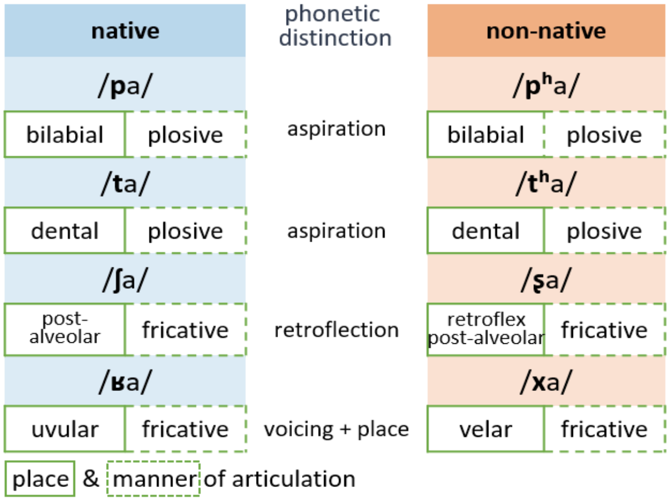


**Figure S8**. Syllable stimuli and phonetic features of the consonants. The native tokens (/pa/, /ta/, /ʃa/ and /ʁa/) are in blue, while the non-native Mandarin tokens (/pʰa/, /tʰa/, /ʂa/ and /xa/) are in orange. For each language, rectangles with green solid lines indicate the place of articulation of the consonants, those with green dashed lines indicate their manner of articulation. The phonetic differences between the native and non-native consonants in each pair are summarized in the middle.

**Table S3.** Acoustic measurements of the consonants embedded in the syllable stimuli. We measured voice-onset-time (VOT) for the plosives and the center-of-gravity (CoG), skewness and kurtosis for the fricatives. VOT is the time interval between the release of a plosive/stop consonant and the onset of vocal fold vibration during the production of the subsequent vowel. CoG represents the spectral mean frequency of a fricative, reflecting where the sound energy is concentrated. Skewness indicates the asymmetry of the spectral energy distribution of fricatives, and kurtosis measures the peak of the spectral distribution.

| **Plosives** | VOT (ms) | **Fricatives** | CoG (Hz) | Skewness | Kurtosis |
|---|---|---|---|---|---|
| **p** | 25.80 | **ʃ** | 4739.18 | 1.17 | 1.70 |
| **pʰ** | 108.26 | **ʂ** | 3459.95 | 1.86 | 3.40 |
| **t** | 43.60 | **ʁ** | 805.16 | 6.02 | 50.23 |
| **tʰ** | 113.40 | **x** | 1373.22 | 5.03 | 36.79 |

**Behavioral task - Familiarization and screening**

All participants first performed a behavioral inclusion test to ensure that their individual level of performance fit the inclusion criteria for the fMRI experiment. The task was a two-alternative forced choice (2AFC) task in which participants had to categorize the heard consonants, embedded in consonant-vowel (CV) syllables, as native or foreign speech sounds. To first familiarize them with the task and the perceptual differences between native and non-native consonants, participants listened to each pair of intact syllables twice (i.e. /pa/ vs. /pʰa/, /ta/ vs. /tʰa/, /ʃa/ vs. /ʂa/, and /ʁa/ vs. /xa/). During the stimulus presentation, native consonants were illustrated on the computer screen by a French flag and non-native consonants by an earth icon. Participants then listened to each pair of intact syllables, which were presented twice in total but in a reversed order (e.g., /pa-pʰa/ or /pʰa-pa/) without any simultaneous icon in the screen. They had to categorize the second consonant as native or as a sound from a foreign language. To give their response, they had to press one of two buttons assigned to the corresponding icon on the screen with their left index or middle finger. The orders of syllable pairs and the icons were randomized across participants. Feedback was given after each trial by either a green circle framing the correct response (i.e. correct icon), or a red circle framing the expected correct answer. After this familiarization, the inclusion task then started. Participants first underwent a practice block featuring 8 trials with intact CV triplets of each consonant (e.g., /pa pa pa/) presented in a randomized order. They performed the 2AFC categorization task (i.e. answering the question whether the heard consonant is French or from a foreign language) by pressing one of the buttons (associated with the two icons on the screen) as accurately and rapidly as possible within 1.5 s. Feedback on both correct and incorrect answers were provided, and a written message was displayed if the response was not given within the allowed time. The intertrial interval (ITI; time interval between the offset of a trial and the onset of the next one, where one trial = fixation point 500 ms + stimulus 1500 ms + response window 1500 ms + feedback 2 s) was 550 ms. The inclusion task *per se* followed exactly the same procedure for two blocks of 48 trials (8 triplets of intact syllables × 6 repetitions) but without any written question nor feedback. The order of the stimuli was randomized, and the association between the button press (finger used) and the icons on the screen was counterbalanced across participants.

Participants who passed this behavioral inclusion task (75% and 70% of correctly recognized native and non-native consonants, respectively) then underwent the fMRI session on a different day. Inside the scanner, a similar familiarization procedure was conducted. The perceptual differences between native and non-native consonants in each pair were presented once as previously described, before two practice blocks for the 2AFC task. The first practice block was similar to the one in the inclusion task with 8 intact triplets. The second block was slightly different as half of the 8 trials (randomly chosen, 2 triplets for each language) were masked by noise. The proper 2AFC task of the fMRI experiment then started as described in the main text (see *Task and experimental procedure*).

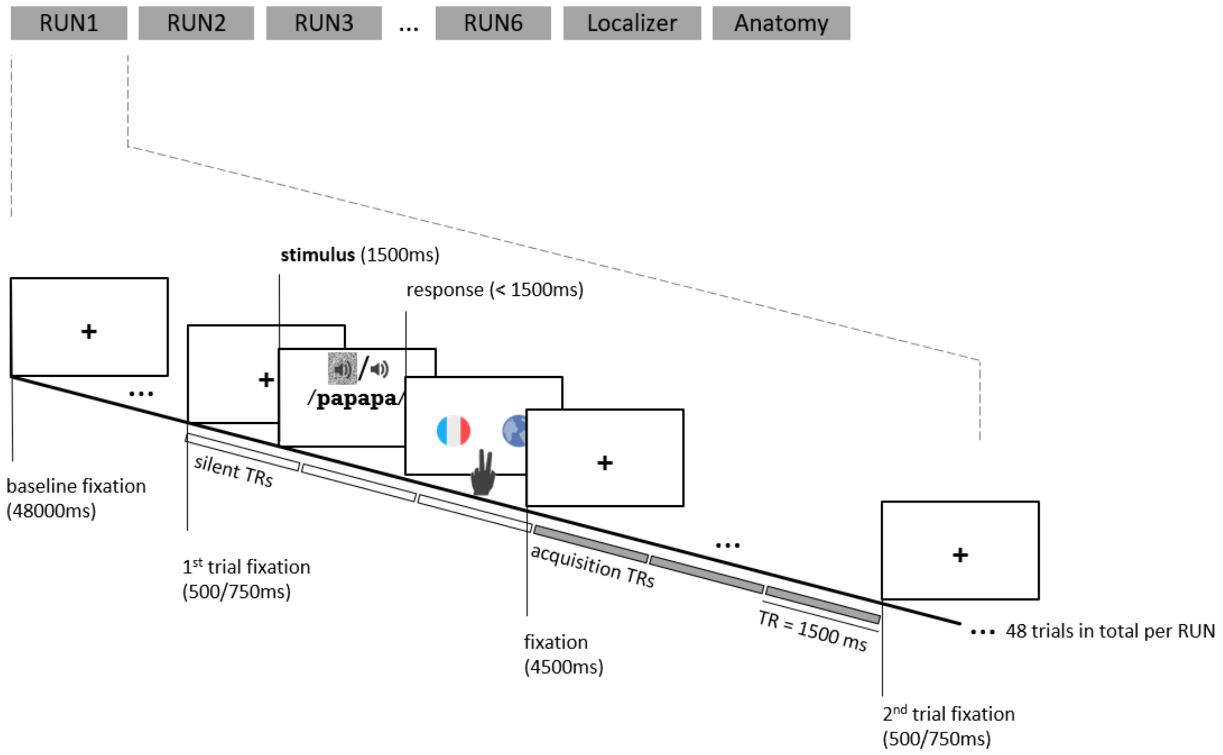


**Figure S9**. Experimental procedure for the 2AFC categorization task on native and non-native consonants and corresponding fMRI acquisition sequence. The experimental session included six blocks, corresponding to six functional runs, during which stimuli (triplets of CV syllables) were auditorily presented during a silent TR (repetition time) using a sparse imaging sequence. Participants were instructed to categorize the heard consonants either as native (French flag) or foreign (earth icon) speech sounds by pressing one of two buttons. The acquisition ended with one block of motor localizer task, corresponding to one functional run using classic continuous acquisition sequence, and an anatomical run.

**Table S4.** Consonant perception contrasts for each condition versus baseline. To assess the neural activation in each condition, contrasts against the baseline were calculated as indicated in the Table for each language condition under each perceptual condition. These contrasts were further used in our MVPA (see *Multivariate Pattern Analysis* in the main text).

**Univariate analyses - Consonant Perception Contrasts**

| | |
|---|---|
| Intact Native | |
| | Intact Native Bilabials = [intact /p/ - Baseline],<br>Intact Native Dentals = [intact /t/ - Baseline],<br>Intact Native Coronal Fricatives = [intact /ʃ/ - Baseline],<br>Intact Native Dorsal Fricatives = [intact /ʁ/ - Baseline] |
| Noisy Native | |
| | Noisy Native Bilabials = [noisy /p/ - Baseline],<br>Noisy Native Dentals = [noisy /t/ - Baseline],<br>Noisy Native Coronal Fricatives = [noisy /ʃ/ - Baseline],<br>Noisy Native Dorsal Fricatives = [noisy /ʁ/ - Baseline] |
| Intact Non-native | |
| | Intact Non-native Bilabials = [intact /pʰ/ - Baseline],<br>Intact Non-native Dentals = [intact /tʰ/ - Baseline],<br>Intact Non-native Coronal Fricatives = [intact /ʂ/ - Baseline],<br>Intact Non-native Dorsal Fricatives = [intact /x/ - Baseline] |
| Noisy Non-native | |
| | Noisy Non-native Bilabials = [noisy /pʰ/ - Baseline],<br>Noisy Non-native Dentals = [noisy /tʰ/ - Baseline],<br>Noisy Non-native Coronal Fricatives = [noisy /ʂ/ - Baseline],<br>Noisy Non-native Dorsal Fricatives = [noisy /x/ - Baseline] |

**Table S5.** Mouth Motor Localizer contrast. The Mouth Motor Localizer contrast (see *Univariate Analyses*) includes brain activity evoked by the movements of both articulators of interest (lips + tongue) contrasted with those of the left and right fingers, to identify articulatory representations for cross-modal decoding. The Table indicates the individual peak coordinates of the decoding center in each hemisphere (see proximate visualized loci in Figure S10).

**fMRI MVPA - Cross-modal decoding**

| Participants | Left hemisphere | Right hemisphere |
|---|---|---|
| P1 | [-64, 2, 13] | [57, -6, 40] |
| P2 | [-48, -14, 51] | [52, -14, 51] |
| P3 | [-67, -6, 13] | [49, -9, 38] |
| P4 | [-56, -6, 32] | [60, 5, 30] |
| P5 | [-56, -6, 38] | [54, -11, 48] |
| P6 | [-45, -11, 35] | [49, -6, 38] |
| P7 | [-56, -11, 43] | [60, -6, 38] |
| P8 | [-56, -6, 48] | [52, -9, 46] |
| P9 | [-64, -3, 21] | [60, -1, 24] |
| P10 | [-51, -11, 30] | [52, -9, 27] |
| P11 | [-43, -11, 46] | [43, -11, 46] |
| P12 | [-56, -3, 48] | [57, 10, 43] |
| P13 | [-56, -11, 43] | [57, -6, 35] |
| P14 | [-43, -17, 40] | [49, -11, 43] |
| P15 | [-59, -14, 46] | [60, -9, 43] |
| P16 | [-56, -3, 43] | [46, -11, 38] |
| P17 | [-45, -9, 46] | [49, -11, 46] |
| P18 | [-43, -14, 38] | [60, -6, 43] |
| P19 | [-51, -14, 43] | [52, -14, 40] |
| P20 | [-51, 2, 57] | [60, -3, 35] |
| P21 | [-48, -10, 40] | [49, -8, 37] |
| P22 | [-56, -6, 40] | [54, -6, 40] |
| P23 | [-51, -14, 46] | [52, -9, 48] |
| P24 | [-53, -11, 40] | [62, -9, 38] |

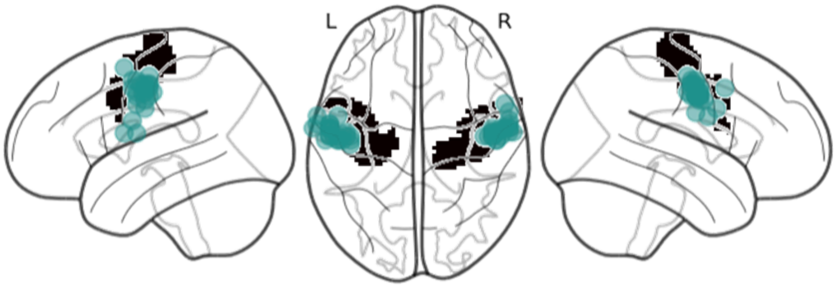


**Figure S10**. Individual peak coordinates visualized in each hemisphere for the Mouth Motor Localizer contrast used in the cross-modal decoding analysis. Each individual peak coordinate defined the center of a 10mm spherical radius, serving as the decoding center for both the movement and perception data (see *Cross-Modal Classification* in Methods). The visualized spherical size for each participant (green areas) does not represent the actual 10mm radius in each individual brain. Black areas indicate the location of the precentral gyrus defined by the AAL atlas (Tzourio-Mazoyer et al., 2002) in a template glass brain using Python Nilearn (Abraham et al., 2014).
